# An IMPDH2 variant associated with neurodevelopmental disorder disrupts purine biosynthesis and somitogenesis

**DOI:** 10.1101/2025.05.09.652712

**Authors:** Audrey G. O’Neill, Morgan E. McCartney, Gavin M. Wheeler, Jeet H. Patel, Gardenia Sanchez-Ramirez, Justin M. Kollman, Andrea E. Wills

## Abstract

IMP dehydrogenase (IMPDH) controls a key regulatory node in purine biosynthesis. Gain-of-function mutations in human IMPDH2 are associated with neurodevelopmental disorders and neuromuscular symptoms including dystonia, but the developmental mechanisms underlying these defects are unknown. We previously showed that these mutants are insensitive to GTP inhibition and hypothesized that their hyperactivity would affect nucleotide metabolism *in vivo*. Here, we characterize the metabolic and developmental consequences of the neurodevelopmental disorder-associated IMPDH2 mutant, S160del, in *Xenopus tropicalis*. We show that expressing S160del but not WT human IMPDH2 disrupts purine pools and somitogenesis in the developing tadpole. We also show that S160del disrupts *in vivo* IMPDH filament assembly, a well-described IMPDH regulatory mechanism. Cryo-EM structures show that S160del disrupts filament assembly by destabilizing the dimerization of regulatory Bateman domains. Dimerization of Bateman domains and subsequent filament formation can be restored with a high affinity ligand, but this does not restore sensitivity to GTP inhibition, suggesting S160del also disrupts allostery of IMPDH2 filaments. This work demonstrates that the structural effects of patient IMPDH2 variants can cause disruptions both to nucleotide levels and to the normal development of sensorimotor structures, helping us better understand the physiological basis of disease in these patients.

**SIGNIFICANCE STATEMENT:** IMPDH2 is a critical enzyme for *de novo* purine biosynthesis, regulating the balance between adenine and guanine nucleotides. Under purine stress, it forms filaments that resist feedback inhibition by GTP. Patients with gain-of-function variants of this enzyme suffer from early-onset neuromotor symptoms including dystonia. Here, we express one gain-of-function variant of IMPDH2, S160del, in *Xenopus tropicalis*. S160del is particularly powerful for structural and developmental studies, as it impedes filament formation and also is insensitive to feedback inhibition by GTP. Here, we show S160del can perturb vertebrate development, metabolism, and filament formation in a dominant fashion. Insights from this work will open the door to a new suite of studies defining the function of purine metabolism in development and disease.

## INTRODUCTION

Purine nucleotides are essential building blocks of life. Beyond being critical components of DNA and RNA, they act as cellular currency, signaling molecules, second messengers, and allosteric effectors. They are also essential for cytoskeletal function and remodeling. To maintain the appropriate levels of purine nucleotides in the cell, there are two biosynthetic pathways to produce them – the salvage pathway and the *de novo* pathway. Mutations in enzymes of both pathways have been linked to a range of neuromuscular and neurodevelopmental disorders (1, 2). Of these enzymes, IMP dehydrogenase (IMPDH), which catalyzes the first committed step of guanine nucleotide biosynthesis, controls an important regulatory node in the *de novo* pathway, and is regulated through allosteric inhibition by GTP. Additionally, the assembly of IMPDH into micron-scale filament structures tunes this regulation by reducing sensitivity to GTP inhibition (3, 4).

There are two isoforms of IMPDH in humans, and several lines of evidence implicate IMPDH2 as the critical isoform in early development. IMPDH2 is highly expressed across most tissues and is specifically upregulated in proliferative cells and in early development (5, 6). In zebrafish, IMPDH2, but not IMPDH1, is predominantly expressed in the somites (7). While IMPDH1 expression is specifically higher than IMPDH2 in the developed retina, only IMPDH2 is detectable in early retinal development (5, 8). In 2020, a cohort of individuals heterozygous for *de novo* missense mutations in IMPDH2 were described to exhibit a range of neurodevelopmental phenotypes (9). While patient symptoms varied widely, more common symptoms included hypotonia, developmental delay, intellectual disability, and dystonia, a hyperkinetic movement disorder. Recently, we showed through *in vitro* biochemical assays that the IMPDH2 variants expressed in these patients are insensitive to feedback inhibition by the downstream product GTP, and we characterized these mutations as gain-of-function (10). Some mutations also affected the ability of IMPDH2 to assemble into filamentous oligomers, which are necessary for allosteric regulation (3, 4). IMPDH filaments also appear to play a role in the development of mice, though its specific early developmental functions are not fully characterized (11). Collectively, these features suggest that IMPDH2 function and filament assembly may be crucial for the functional development of neural or muscle tissues, motivating our study.

While our *in vitro* characterization of IMPDH2 mutants suggested that metabolism and the developmental processes that rely on purine nucleotides would be affected by IMPDH2 dysregulation, an animal model is needed to uncover molecular mechanisms of disease. *Xenopus laevis* and *Xenopus tropicalis* are widely used to model human disease (12–14). Their transparent tadpoles also facilitate *in vivo* and fixed imaging, including of IMPDH filaments that are induced by treatment with the IMPDH inhibitor mycophenolic acid (MPA), as we previously showed (15). Recently, loss of function studies of other purine biosynthetic enzymes in *Xenopus laevis* revealed that these enzymes play required roles in somitogenesis, with their loss resulting in phenotypes that resemble the hypotonia documented in patients (16). Taken together, *Xenopus* is a useful model for studying purine biosynthesis in development.

Here, we exploited the specific structural properties of the patient variant S160del to better understand the function of IMPDH2 in embryonic development, using *Xenopus tropicalis* as a model. The in-frame deletion of serine 160 (S160del) is associated with neurodevelopmental disorders in patients (9). We previously showed using negative stain electron microscopy (EM) on recombinantly expressed and purified protein that the S160del variant does not assemble into filaments *in vitro,* and also resists feedback inhibition by GTP (10). These observations led us to the hypothesis that S160del may perturb physiological levels of purine metabolites in developing tissue, resulting in developmental defects. Here, we confirm this, finding that expression of hIMPDH2-S160del in *Xenopus tropicalis* disrupts purine biosynthesis and somitogenesis in the developing tadpole. Additionally, S160del expression prevents filament formation by endogenous IMPDH in tadpoles, supporting our previous *in vitro* results. To understand how S160del disrupts filament formation, we demonstrated with cryo-EM that the deletion of S160 prevents dimerization of the regulatory Bateman domain. To test if stabilization of the Bateman dimer interface would restore sensitivity to feedback inhibition, we showed that the dinucleoside polyphosphate, Ap5G, promotes S160del filament formation. However, these S160del filaments are insensitive to GTP inhibition, suggesting that filament formation and dysregulation are separable features in this mutant. Because the deletion of S160 does not affect the structure of the catalytic domain, inhibitors targeting the active site, such as MPA may be a useful therapeutic strategy. Taken together, this work introduces a model for characterizing the mechanisms of disease resulting from a gain-of-function IMPDH2 mutant. Our work demonstrates that the structural changes introduced by S160del are coupled to dominant perturbations of cell metabolism and neurodevelopment. This establishes a framework for understanding how other variants of purine biosynthetic enzymes may also lead to developmental defects.

## RESULTS

### S160del disrupts purine nucleotide levels in *Xenopus tropicalis* tadpoles

Because patients were heterozygous for these gain-of-function IMPDH2 variants, we hypothesized that the mutants would exert a dominant effect on early development (9, 10). We therefore sought to interrogate the effects of adding S160del to the endogenous functions of IMPDH2 *in vivo*. To test this, we overexpressed WT and S160del human IMPDH2 (hIMPDH2) in *Xenopus tropicalis* embryos, which share 93% amino acid sequence identity with hIMPDH2 (Supplemental Fig. 1). We synthesized mRNA transcripts encoding WT or S160del hIMPDH2 and injected this mRNA into both cells at the 2-cell stage. Western blots of whole embryo lysates at stage 21, approximately 1 day post fertilization (dpf), confirmed dose-dependent overexpression of the injected mRNA compared to uninjected controls (Fig. 1D, Supplemental Figs. 2-3). We detected the persistence of overexpressed IMPDH2 via western blot up to stage 47, approximately 5 dpf, for the 1000 pg dose of mRNA (Supplemental Fig. 4).

**Figure 1.**
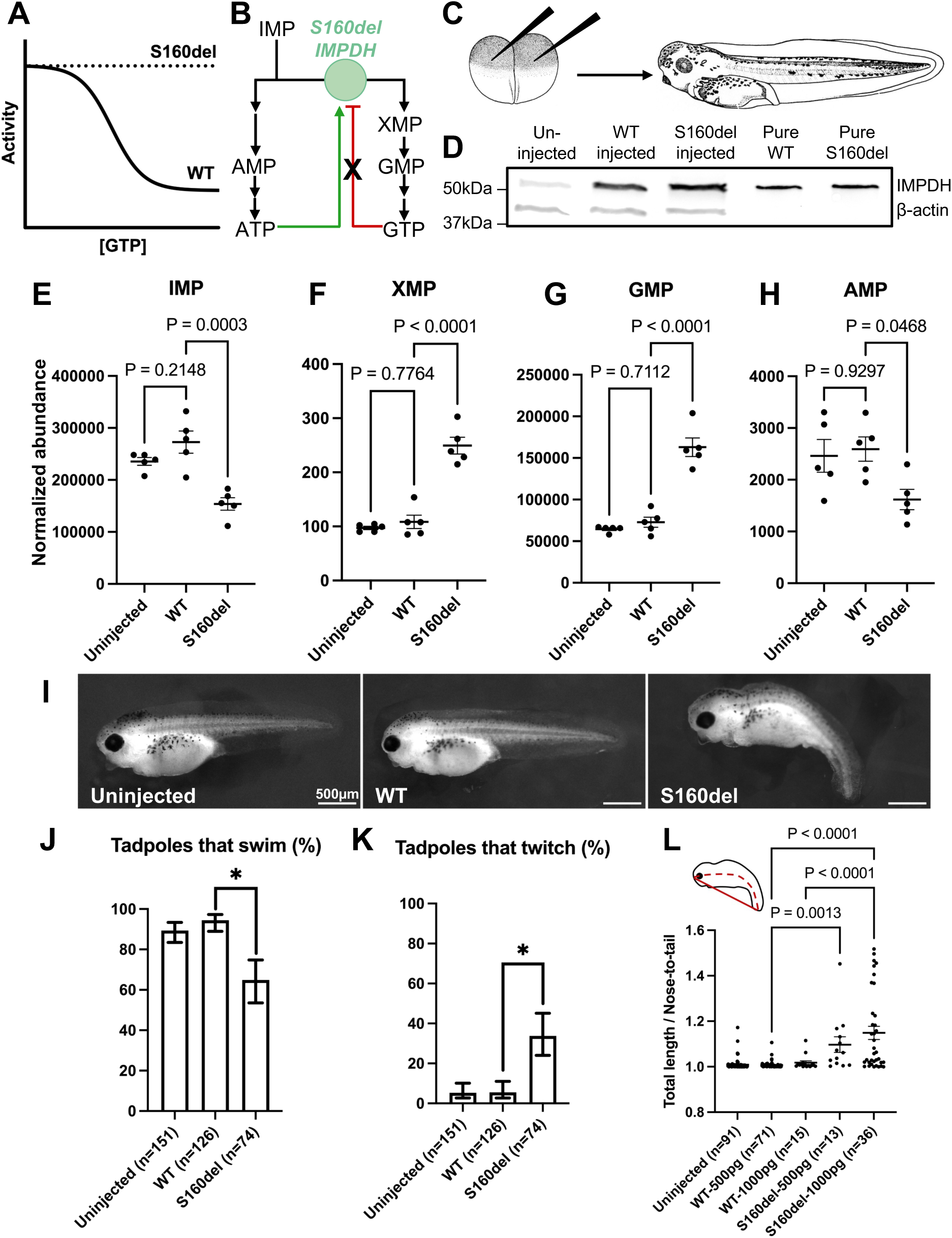
**Overexpression of human IMPDH2 in *Xenopus tropicalis*** A. Representative plot of IMPDH2 activity under increasing concentrations of GTP for WT and S160del hIMPDH2. B. Schematic diagram of *de novo* purine synthesis. The S160del mutation disrupts normal allosteric feedback inhibition by GTP, represented as a black (X) in the diagram. C. Experimental diagram. *Xenopus tropicalis* embryos are injected with mRNA into each cell at the 2-cell stage and then raised to NF stage 41 for phenotype and metabolic assay. Drawings from Nieuwkoop and Faber (1994) Normal Table of Xenopus laevis (Daudin). Garland Publishing Inc, New York ISBN 0-8153-1896-0. Copyright © 1994 Pieter D. Nieuwkoop and J. Faber. Digital images created by and accessed from XenBase. D. Representative western blot analysis of overexpression of hIMPDH2 in *Xenopus tropicalis* embryos at NF stage 21, following mRNA injection. E-H. Quantification of selected purine synthesis metabolites from LC-MS assay. Each data point represents normalized metabolite abundance of an aggregate tissue sample composed of 10 whole tadpoles. One-way ANOVA and Tukey’s multiple comparisons test performed on metabolite levels normalized to total protein from BCA. Plotting mean +/-SEM. I. Photographs of NF stage 41 tadpoles following injection with the indicated mRNA. J and K. Quantification of behavior effects of IMPDH2 overexpression. Plotting percentage +/- the 95% confidence interval of the proportion, calculated using the Wilson/Brown method. Chi-square analysis indicated significance. The Marasculio procedure was used to determine if WT and S160del groups were significantly different, at P=0.05. L. Quantification of ventral tail curvature. One-way ANOVA and Tukey’s multiple comparisons test. Plotting mean +/- SEM.

We then investigated the mutant’s effect on metabolism in the developing tadpole through liquid chromatography-mass spectrometry (LC-MS) of whole tadpole lysates, collected at stage 41, approximately 3 dpf. Our previous *in vitro* characterization of S160del led us to propose that its insensitivity to feedback inhibition would result in a gain-of-function effect *in vivo* (10). We hypothesized that expression of this mutant would result in increased pools of guanine nucleotides over adenine nucleotides in purine biosynthesis. Strikingly, we found that while WT-expressing tadpoles had no significant difference in metabolite levels compared to uninjected tadpoles, the metabolic profile of S160del-expressing tadpoles had significantly shifted (Supplemental Fig. 6, Supplemental Fig. 7). The downstream metabolites XMP, GMP, guanine, guanosine, and xanthosine were significantly elevated (Fig. 1F-G, Supplemental Fig. 8). Conversely, IMP and AMP levels were decreased in the S160del group (Fig. 1E, 1H).

The S160del group also had elevated levels of urate, a byproduct of purine nucleotide degradation (Supplemental Fig. 8). These findings demonstrate that S160del specifically disrupts the physiological abundance of purine nucleotides, supporting the hypothesis that the gain-of-function effect of S160del impacts relevant metabolite levels.

### S160del expression affects early *Xenopus tropicalis* development

After establishing that S160del expression affects metabolism in an animal model, we sought to understand its impact on early development. We began with behavioral and morphological characterization of tadpoles expressing S160del. At stage 41, tadpoles expressing WT or S160del hIMPDH2, as well as uninjected controls, were tested for their ability to swim away from physical stimulus with a blinded escape reflex assay (17). Tadpoles expressing S160del were significantly less responsive than the dose-matched WT groups and the uninjected controls (Fig. 1J). They also twitched spontaneously, without any external stimulus (Fig. 1K, Supplemental Movies). Following behavioral scoring, these same tadpoles were fixed and imaged on a stereomicroscope for morphological phenotyping and measurements of tail curvature (18). Both groups expressing S160del were significantly shorter (Supplemental Fig. 5) and had significantly more ventrally curved tails, compared to the dose-matched WT groups and the uninjected controls (Fig. 1I, 1L). Notably, tadpoles injected with 1000 pg of hIMPDH2-WT mRNA were also shorter than the uninjected controls (Supplemental Fig. 5). However, the ventral tail curvature and the twitching behavior cannot be attributed to overexpression of hIMPDH2, as the controls overexpressing wildtype hIMPDH2 at similar levels were indistinguishable from uninjected controls with endogenous IMPDH2 levels. These phenotypes resemble patient symptoms of hyperkinetic movements and abnormal posturing and suggest that expression of S160del in a vertebrate system specifically results in morphological and neurobehavioral defects during development.

### Somite boundaries are disorganized in tadpoles expressing S160del

The motor defects and tail morphology in S160del-expressing tadpoles led us to hypothesize that there were underlying defects in either muscle or neuronal development. To assess tissue-specific defects in the developing tadpoles, we performed immunostaining of neurofilament and skeletal muscle on fixed tadpoles at stage 41. From the neurofilament staining, tadpoles expressing S160del had a significantly lower density of intersomitic axon bundles along the length of the tail, and organization of the remaining axons was disrupted (Fig. 2B, 2D). Notably, the WT-injected group had a slightly higher number of axon bundles per millimeter as compared to the uninjected controls (Fig. 2D). From the skeletal muscle staining, there was also a clear defect in the number and clarity of somite boundaries, with poorly defined or indistinguishable somite boundaries in tadpoles expressing S160del (Fig. 2C, 2E). Taken together, these results suggest a defect in myogenesis and neurofilament distribution resulting from S160del expression, resembling the hypotonia of patients.

**Figure 2.**
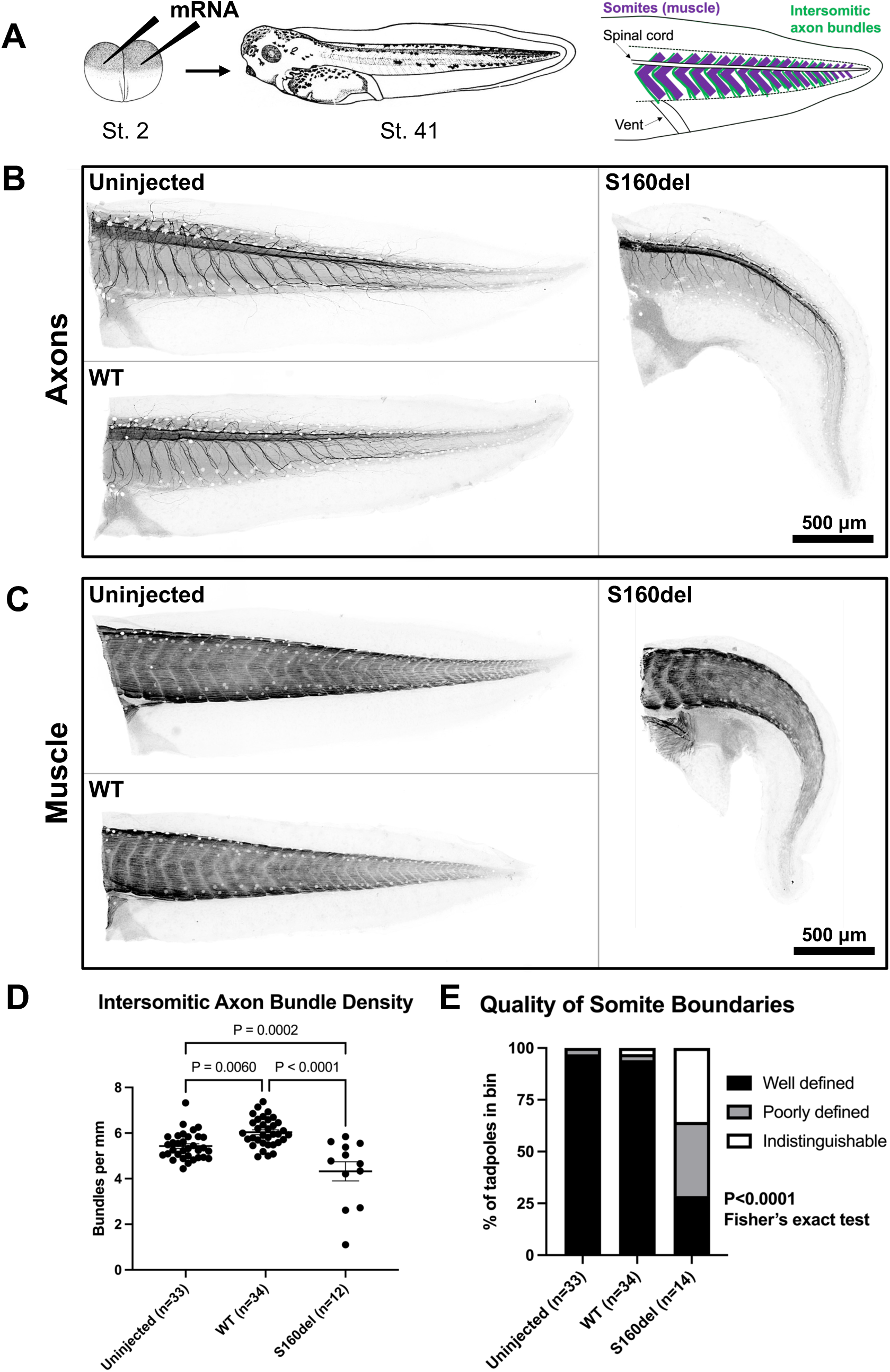
**Tadpoles expressing hIMPDH2-S160del have somitic defects** A. Diagrams of experimental design and simplified *Xenopus tropicalis* tail anatomy. *Xenopus tropicalis* embryos are injected with 500 pg of mRNA into each cell at the 2-cell stage and then raised to NF stage 41, approximately 3 dpf. B. Representative anti-neurofilament immunofluorescence images of tadpoles injected with the indicated mRNA. C. Representative anti-skeletal muscle immunofluorescence images of tadpoles injected with the indicated mRNA. D. Quantification of the number of intersomitic axon bundles per mm in the tail. One-way ANOVA and Tukey’s multiple comparisons test. E. Quantification of somite boundary quality. Fisher’s exact test for categorical variables.

### S160del has a dominant negative effect on IMPDH assembly *in vivo*

To understand how S160del might exert a dominant effect, we sought to determine whether it might disrupt one of the key properties of IMPDH2: its ability to form rod and ring filament structures. We previously described the formation of rods and rings in the developing tadpole under metabolic challenges, including tail regeneration and treatment with the IMPDH inhibitor mycophenolic acid (MPA) (15). In wildtype, uninjected tadpoles, IMPDH2 expression was primarily diffuse in the axial skeletal muscle when observed via immunostaining (Fig. 3E). However, upon MPA treatment, we observed robust rod and ring formation from endogenous IMPDH2, particularly along the boundaries of the somitic muscles (Fig. 3B). Similarly, tadpoles expressing WT hIMPDH2 formed rods and rings in response to MPA treatment. Notably, even without MPA treatment, some rods and rings were present in WT hIMPDH2-expressing tadpoles, presumably due to the effects of overexpression (Fig. 3C,F). Overexpression of IMPDH has been reported to induce in rods and rings in previous cell culture studies (8, 19, 20).

**Figure 3.**
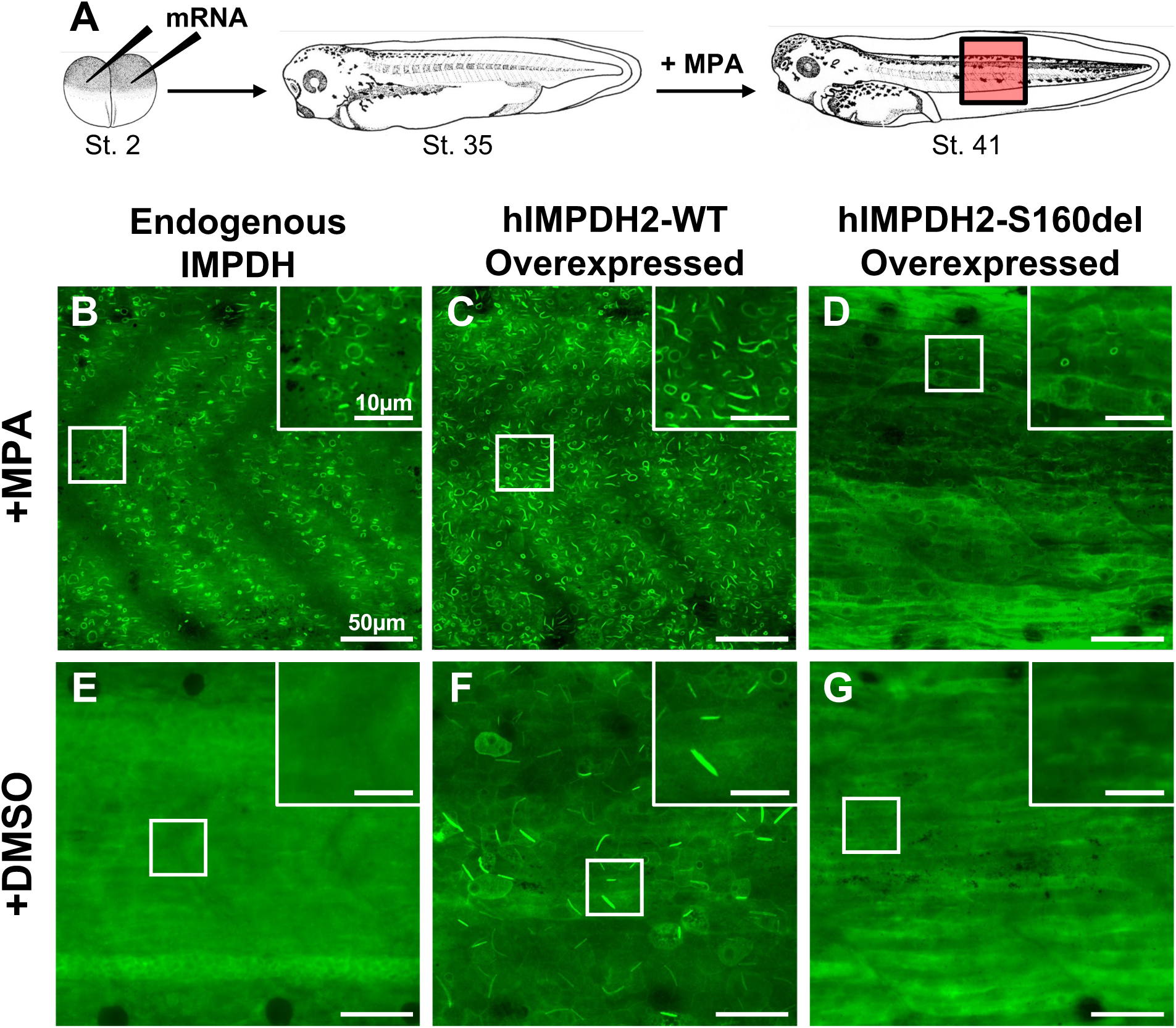
**S160del destabilizes IMPDH superstructures *in vivo*** A. Experimental diagram. *Xenopus tropicalis* embryos are injected with 500 pg of mRNA into each cell at the 2-cell stage, raised to NF stage 35 (approximately 2 dpf), and then treated with either 1 uM MPA or vehicle control. Tadpoles are raised for an additional 24 hours to NF stage 41 and then stained for IMPDH2. B-G. Representative anti-IMPDH2 immunofluorescence images of uninjected (endogenous) or mRNA injected tadpoles treated with either 1 uM MPA (B-D) or DMSO vehicle control (E-G). Insets in the top right corners of each image are close-ups of the regions indicated by white boxes.

In contrast to uninjected and WT-injected tadpoles, tadpoles expressing S160del hIMPDH2 exhibited a dominant-negative loss of rod and ring formation under MPA treatment, with fewer structures observed (Fig. 3D,G). This data indicates that the structural phenotype we have observed *in vitro* for S160del translates to a defect in rod and ring formation in the cells of a vertebrate system. S160del is likely able to co-assemble with endogenous IMPDH into heterooligomers, likely preventing endogenous IMPDH filament assembly.

### S160del disrupts the dimerization of Bateman domains in human IMPDH2

After demonstrating that S160del expression has a dominant negative effect on IMPDH filament assembly in *Xenopus tropicalis*, we wanted to characterize its disruption of filament assembly at higher resolution. We previously showed with negative stain EM that the S160del mutant is unable to form filaments following incubation with ATP or GTP (10). S160 is located between allosteric sites 1 and 2, and forms hydrogen bonds with the phosphates of ATP in site 1 (Fig. 4A). Additionally, several residues downstream of S160 are involved in Bateman dimer interface contacts (Supplemental Fig. 9). We wondered if the deletion of S160 would weaken nucleotide binding in allosteric site 1, or if its deletion would result in misfolding of the Bateman domain.

**Figure 4.**
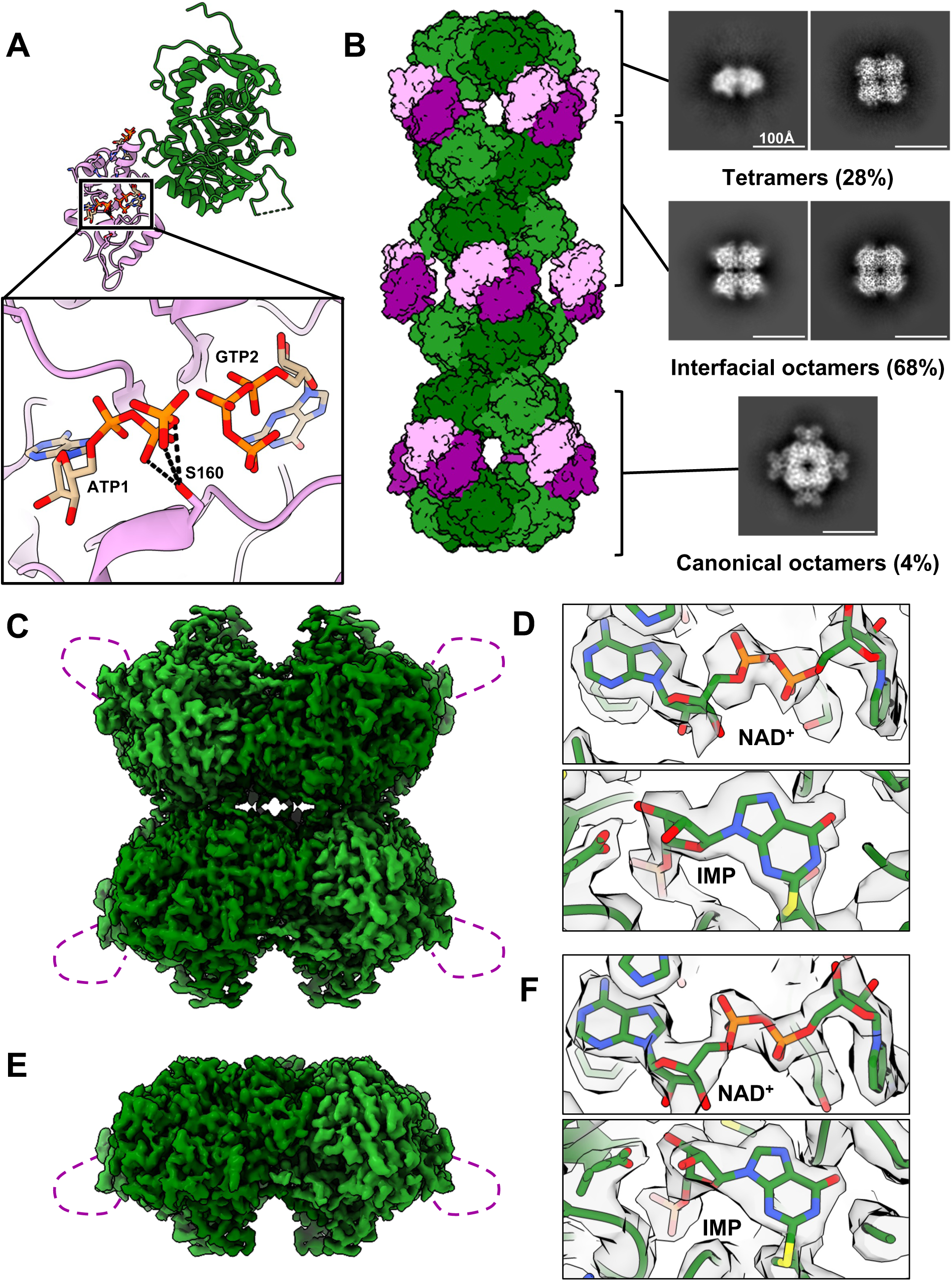
**S160del destabilizes IMPDH2 polymerization *in vitro*** A. Structure of the hIMPDH2 monomer under physiologically relevant ligand conditions (with bound ATP, GTP, IMP, NAD^+^). Serine 160 is located within the regulatory Bateman domain and is proximal to both ATP in site 1 and GTP in site 2 when bound. Potential hydrogen bonds to the phosphates of ATP in site 1 are displayed as dashed lines. B. Distribution of S160del assembly states as determined by cryo-EM. Selected 2D class averages are shown. The majority of selected particles (68%) are classified as interfacial octamers with a smaller subset (28%) of tetramers. Only 4% of selected particles existed in the canonical octamer state, but secondary structure is apparent in the Bateman domains. C-D. Cryo-EM map of the S160del interfacial octamer (C). Only the catalytic domain is resolved (green). The disordered Bateman domain is represented as a purple dashed line. Both NAD^+^ and IMP are bound (D). E-F. Cryo-EM map of the S160del tetramer (E). Only the catalytic domain is resolved (green). The disordered Bateman domain is represented as a purple dashed line. Both NAD^+^ and IMP are bound (F).

To test this, we collected a cryo-EM dataset of S160del in the presence of saturating GTP, ATP, IMP, and NAD+. As was observed with negative stain, no filaments were present. The dataset was compositionally heterogeneous, with 28% of the selected particles classified as tetramers and 68% of the selected particles classified as an interfacial octameric assembly, in which the filament assembly interface was formed but the Bateman domains were not dimerized (Fig. 4B, Supplemental Fig. 10). The tetramer refined to a global resolution of 2.3 Å with C4 symmetry imposed, and the interfacial octamer refined to 2.1 Å with D4 symmetry imposed (Supplemental Figs. 10-11). In both the tetramer and interfacial octamer reconstructions, the Bateman domains were not resolved, suggesting that they were highly flexible as expected (4) (Fig. 4C, 4E).

Substrates were clearly resolved in the active site in both reconstructions (Fig. 4D, 4F). The catalytic domain of the interfacial octamer reconstruction was nearly identical to that of WT hIMPDH2, with a C*α* RMSD of 0.6 Å (Supplemental Fig. 12A). The tetramer adopted a more bowed confirmation as compared to the interfacial octamer (Supplemental Fig. 12B). Overall, the catalytic domain of the tetramer and interfacial octamer reconstructions was unremarkable as compared to the WT enzyme under the same conditions.

Surprisingly, 4% of the selected particles (31,924 particles) were classified as canonical octamers, with Bateman dimers clearly resolved in the 2D class averages (Fig. 4B, Supplemental Fig. 10). These particles adopted a preferred orientation in ice as has been previously observed for canonical octamers of IMPDH (4), impeding high-resolution structural characterization. However, the presence of the canonical octamer in this dataset, and the secondary structure observed in the 2D class averages, suggest that while S160del clearly disrupts the stability of the Bateman dimer interface, possibly by weakening the binding of nucleotides in allosteric site 1, it does not cause misfolding of the Bateman domain.

### Filament assembly and GTP insensitivity are separable features of S160del

The syndromic effects of the S160 deletion in patients might derive either from its preventative effect on oligomerization, or from its insensitivity to GTP feedback inhibition. Our structural analyses provided a path to test if oligomerization could be restored in this mutant, and whether that would restore GTP sensitivity. According to the current model of IMPDH inhibition, GTP-induced compression of canonical octamers forces the interaction of opposing finger domains (Supplemental Fig. 13). This finger domain interaction inhibits enzyme function by impeding the dynamics of the active site during the catalytic cycle (6, 21–23). In *Mycobacterium smegmatis* IMPDH, compression of the octamer induced by GTP binding causes the finger domain and catalytic flap to adopt a conformation which prevents IMP binding (24). Finger domains would not be positioned to interact at all in the free tetramers or interfacial octamers observed in our cryo-EM dataset of S160del. However, the small population of canonical octamers that we observed in this dataset suggests that S160del is capable of forming the Bateman dimer interface in the presence of nucleotides. We therefore sought to stabilize the formation of S160del filaments and test whether the effect of the S160 deletion on GTP inhibition is dependent on its effect on filament formation.

We hypothesized that we could promote the dimerization of Bateman domains and favor canonical octamer formation by incubating S160del with higher affinity ligands. Dinucleoside polyphosphates have been reported to bind across allosteric sites 1 and 2 of IMPDH with 100x higher affinity than ATP or GTP alone (25). With negative stain EM, we found that S160del was able to polymerize following incubation with Ap5G, confirming that nucleotide binding to allosteric sites 1 and 2 and the subsequent dimerization of Bateman domains is not completely disrupted by the deletion of S160 (Fig. 5B). Interestingly, these filaments remained insensitive to GTP inhibition, suggesting that the effect of S160del on filament formation is independent from its effect on allosteric regulation (Fig. 5C). These results provide a new strategy for isolating these properties in other disease-associated IMPDH2 mutants that destabilize filament formation. Additionally, they inform drug-design approaches for treating disorders that result from IMPDH2 dysregulation.

**Figure 5.**
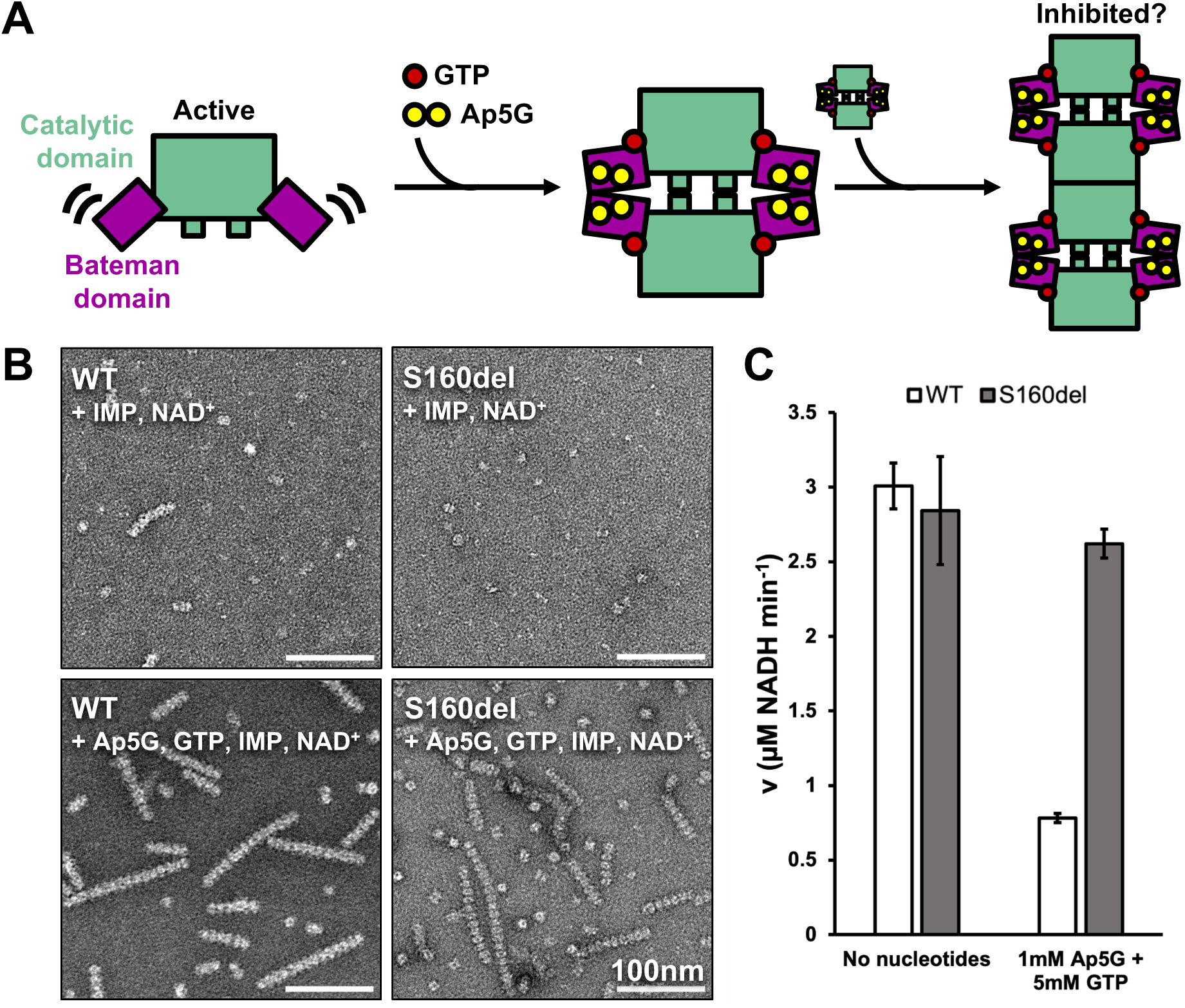
**Ligand Binding restores polymerization, but not GTP regulation, to S160del** A. Experimental diagram. S160del IMPDH2 tetramers are active, and the regulatory Bateman domain (purple) is highly flexible. Ap5G and GTP bind to the Bateman domains of tetramers and stabilize the formation of canonical octamers and filaments. B. Negative stain electron micrographs of WT and S160del hIMPDH2 in the assay conditions from panel C. C. *In vitro* enzyme assay of WT and S160del hIMPDH2 in the absence (left) or presence (right) of Ap5G and GTP. Plotting mean of three technical replicates. Error bars represent SD.

## DISCUSSION

Errors in purine and pyrimidine metabolism result in a variety of disorders affecting neuromuscular development, including dystonia (2, 26). Both gain-of-function and loss-of-function mutations have been documented in purine-related developmental disorders, sometimes affecting the same enzyme. Such is the case for phosphoribosylpyrophosphate synthetase (PRPS), a regulatory enzyme upstream of IMPDH in purine biosynthesis, where mutations that decrease activity lead to hearing and vision loss, muscle weakness, and intellectual disability; mutations that increase PRPS activity lead to gout and progressive renal failure. The more severe, early-onset phenotype of PRPS superactivity also includes symptoms similar to those resulting from PRPS deficiency, such as hypotonia and developmental delay (2). This suggests that a delicate balance between metabolites in these pathways is essential for proper functioning of nerves and muscle, and disrupting that balance in either direction can have deleterious consequences on neuromuscular development, sometimes even leading to overlapping clinical presentations.

The gout caused by PRPS superactivity is a result of overproduction of uric acid, the final degradation product of purine nucleotides (27). Build-up of guanine and hypoxanthine, and subsequent overproduction of uric acid, is characteristic of congenital HPRT deficiency, which results in increased flux through *de novo* purine biosynthesis rather than the salvage pathway (28–30). Metabolites like urate and guanine can therefore be useful biomarkers for diagnosing these disorders.

Recently, *Xenopus laevis* was used as a model to study the role of purine biosynthesis in embryonic development (16). Their study focused on the role of adenylosuccinate lyase (ADSL), an enzyme which functions both up- and down-stream of IMPDH in the *de novo* purine biosynthesis pathway. ADSL deficiency in humans also causes developmental delay, autism, epilepsy, and hyperkinetic movements. Knockdown of ADSL in *Xenopus laevis* leads to defects in somitogenesis and hypaxial muscle formation, likely through disrupting the balance of purine nucleotide pools. Knockdown of other purine biosynthetic enzymes, hypoxanthine phosphoribosyltransferase (HPRT) and phosphoribosyl pyrophosphate amido transferase (PPAT) likewise led to the same myogenic defects in tadpoles, and mirrored the hypotonia seen in patients (16).

Here, we show that expressing the gain-of-function IMPDH2 variant S160del in *Xenopus tropicalis* resulted in significantly elevated levels of GMP, guanine, and urate, supporting our hypothesis that the IMPDH2 hyperactivity we characterized *in vitro* would affect metabolite levels *in vivo*. The expression of this hyperactive variant also resulted in behavioral and morphological defects in the developing tadpole, including ventral tail curvature and twitching. Immunostaining of the tail revealed a disorganization of muscle patterning in the trunk, with poorly defined or indistinguishable somite boundaries and disorganized innervation of axon bundles in between somites. These data suggest that disrupted purine pools may be affecting somitogenesis, and that the hypotonia and hyperkinetic movements similar to those observed in patients can be recapitulated in a *Xenopus* model. Future studies may investigate if these metabolomic signatures and tissue-level phenotypes hold in patients.

In addition, we demonstrated that the S160del mutant, which we previously showed is incapable of forming filaments *in vitro*, has a dominant negative effect on the ability of IMPDH to assemble into rod and ring shaped superstructures *in vivo*. With cryo-EM structures of recombinantly purified S160del, we showed that the catalytic domain of S160del is identical to the WT, as expected. We also showed that while S160del primarily assembles into interfacial octamers and tetramers, it is surprisingly still capable of assembling into the canonical octamer. In order to isolate the effects of S160del on filament formation and GTP inhibition, we showed that we can stabilize the formation of S160del filaments by incubating with the dinucleoside polyphosphate Ap5G, which binds to hIMPDH with high affinity (25). Interestingly, while Ap5G promoted S160del polymerization, these filaments remained insensitive to GTP inhibition, suggesting that the deletion of S160 has an effect on allosteric regulation beyond destabilizing filament formation.

This work establishes a model for studying the consequences of IMPDH2 hyperactivity in a vertebrate system, interrogating the embryonic origin of the disease in parallel with metabolism and enzyme behavior. The gain-of-function S160del variant impairs neuromuscular development and IMPDH filament formation in a dominant fashion and significantly affects purine metabolite levels. Strikingly, overexpression of WT hIMPDH2 did not result in these defects in the developing tadpole, supporting the hypothesis that normal guanine nucleotide levels are sufficient to inhibit the WT enzyme despite overexpression. We conclude that dysregulation of IMPDH2 is likely the main driver of disease in S160del patients, rather than changes in overall IMPDH protein levels. Future work may focus on using this system to test IMPDH inhibitors as potential therapeutics for disorders related to IMPDH2 hyperactivity.

## METHODS

### *X. tropicalis* husbandry and use

Use of *Xenopus tropicalis* was carried out under the approval and oversight of the IACUC committee at the University of Washington, an AAALAC-accredited institution (IACUC protocol number 4374-01). Ovulation of adult *X. tropicalis* and generation of embryos by natural matings were performed according to published methods (31, 32). Fertilized eggs were de-jellied in 3% w/v cysteine in 1/9x modified frog ringer’s solution (MR) for 10–15 minutes. Embryos were reared as previously described (31). Briefly, animals were reared in petri dishes at a density of no more than 2 tadpoles/mL of 1/9x MR at a temperature of 22 °C. Clutch mates were randomly assigned to treatment groups. Staging was assessed by the Nieuwkoop and Faber (1994) staging series. Tadpoles do not begin to feed independently until approximately stage 45 and so were not fed during the course of these experiments.

### Cloning

Plasmids encoding WT and S160del hIMPDH2 were subcloned from pSMT3 bacterial expression plasmids (pJK053_pSMT3_hIMPDH2 and pJK314_pSMT3_hIMPDH2-S160del) (3, 10) into the pCS2+8 vector (33) using Gibson assembly. The pCS2+8 vector was a gift from Amro Hamdoun (Addgene plasmid # 34931 ; http://n2t.net/addgene:34931 ; RRID:Addgene_34931). The IMPDH2 coding region was amplified with the following primers: hIMPDH2 template (forward - TTGTTCTTTTTGCAGGATCCATGGCCGACTACCTGATTAGTG; reverse - ATTAATGGCGCGCCACTAGTTCAGAAAAGCCGCTTCTCATACGAATGGAG) pCS2+8 vector (forward-TGAGAAGCGGCTTTTCTGAACTAGTGGCGCGCCATTAATTAAAG; reverse-CACTAATCAGGTAGTCGGCCATGGATCCTGCAAAAAGAACAAGTAGCTTG). PCR products were gel extracted. Insert and backbone were ligated at 50°C for 1 hour in a Gibson assembly reaction. Gibson products were transformed into chemically competent TOP10 *E. coli* cells and grown on LB agar plates containing 100 µg/ml carbenicillin. Individual colonies were picked and grown for 6 hours at 37°C in LB medium + 100 µg/ml carbenicillin. Plasmid DNA was purified using the GeneJET Plasmid Miniprep Kit (Thermo Fisher Scientific). Final construct sequence was verified with both traditional Sanger sequencing of the insert (Genewiz) and Nanopore sequencing of the whole plasmid (Plasmidsaurus) before using for *in vitro* transcription.

### Preparation of mRNA

Plasmid DNA was linearized by overnight digestion with *NotI* enzyme at 37 °C (New England Biolabs). Linear DNA was purified by phenol:chloroform:isoamyl alcohol (25:24:1 v/v) extraction followed by precipitation in isopropanol with 0.05 M sodium acetate. Transcription of mRNA was performed using mMESSAGE mMACHINE SP6 Transcription Kit (Invitrogen). Template DNA was degraded by treating with DNase for 10 minutes at 37°C. The resulting mRNA product was purified by two rounds of precipitation: first in 2.5 M lithium chloride and second in ethanol with 0.14 M ammonium acetate to remove residual lithium ions. Presence of unfragmented mRNA was verified by agarose gel electrophoresis before resuspending in nuclease-free water to a stock concentration of 500 ng/μL.

### Microinjections

Injection mixtures were prepared using mRNA, nuclease-free water, and fluorescent dextran fluoro-Ruby (Invitrogen) to an mRNA concentration of 250 pg/nL. Needles for injections were created from thin wall glass capillaries containing filament (World Precision Instrument) using a needle puller. Embryos were collected in batches of approximately 50 onto a petri dish coated with a thin layer of 0.1% agarose prepared in 1/9x MR to prevent dehydration. A Picospritzer III (Parker Hannifin Corporation) connected to a micromanipulator (Narishige) was used to deliver 2 nL of injection mixture into each blastomere of the 2-cell stage embryo, for a total of 1000 pg of mRNA per embryo. Injected embryos were moved into a recovery solution of 3% w/v Ficoll PM400 (Cytiva) in 1/9x MR for no more than 2 hours before transferring into 1/9x MR rearing media.

### Western Blotting

Four whole embryos or tadpoles per group were homogenized on ice in 100 μL of lysis buffer (50 mM Tris pH 7.6, 150 mM NaCl, 10 mM EDTA, 0.1% Triton X-100, Roche cOmpleteTM Protease Inhibitor Cocktail). Homogenized samples were centrifuged at 18,400xg for 20 minutes at 4°C. The soluble fraction was collected, and samples were denatured by adding 1/4 volume of 4X Protein Sample Loading Buffer (Licor 928-40004). Samples were heated at 100 °C for 5 minutes. Equal inputs of the samples were run on a 4-20% Mini-PROTEAN TGX precast gel (BIO-RAD 4561094) at 180 V for 40 minutes in manufacturer recommended running buffer (25 mM Tris, 192 mM glycine, 0.1% SDS, pH 8.3). Transfer to a nitrocellulose membrane was done using an Invitrogen Power Blotter Select Transfer Stack (Thermo Fisher PB3310) on an Invitrogen Power Blotter System (Thermo Fisher PB0012) using the Mixed Range MW Pre-Programmed method (constant 2.5 A with 25 V limit for 7 minutes). Membrane was incubated in Intercept PBS Blocking Buffer (Licor 927-70001) for 1 hour at 25°C with rocking. Membrane was added into 5 mL of 1X PBS-Tween (137 mM NaCl, 2.7 mM KCl, 10 mM Na2HPO4, 1.8 mM KH2PO4, 0.1% w/v Tween-20 detergent) and 5 mL of Intercept PBS Blocking Buffer. Membrane was incubated in primary antibodies: 1:1000 rabbit anti-IMPDH2 (Proteintech 12948-1-AP) and 1:3000 mouse anti-β-actin (Santa Cruz Biotech sc-47778) overnight, shaking at 4 °C. Membrane was washed with 1X PBS-Tween for 4 x 5 minutes, then added into 5mL of 1X PBS-Tween mixed with 5mL of Intercept PBS Blocking Buffer. Membrane was incubated in the dark for 1 hour, shaking at 25 °C with secondary antibodies: 1:10000 goat anti-rabbit IgG (H+L) (DyLight 800 Fisher PISA510036) and 1:10000 goat anti-mouse IgG (H+L) (DyLight 680 Fisher PI35518). After 4 x 5 minute washes in 1X PBS-Tween, the membrane was imaged on a LI-COR Imaging System (LI-COR Biosciences).

### Behavioral assays

At stage 41, tadpoles were tested for their ability to swim away from physical stimulus using a blinded escape reflex assay (17). Each tadpole was prodded once with a pipette tip at the tip of their tail, and their ability to swim away was tallied. Tadpoles that appeared to twitch without external stimulus were also tallied. Swimming and twitching were not treated as mutually exclusive behaviors.

### Morphological phenotyping

At stage 41, whole tadpoles were fixed in 1x MEM with 3.7% formaldehyde at 4°C overnight. Tadpoles were imaged in 1x PBS + 0.1% Tween-20 on a bed of 1% agarose using a Leica M205 FA stereomicroscope with a color camera. Image analysis and measurements were done in Fiji.

### Immunostaining

Fixed tadpoles were permeabilized by washing 3 × 20 minutes in 1x PBS + 0.01% Triton X-100 (PBS-Triton). Tadpoles were blocked for 1 hour at room temperature in 10% CAS-block (Invitrogen #00-8120) in PBS-Triton. Tadpoles were then incubated in primary antibody [1:100 rabbit anti-IMPDH2, proteintech 12948-1-AP; 1:50 mouse anti-skeletal muscle marker, DSHB 12/101; 1:50 mouse anti-neurofilament, DSHB 3A10] diluted in 100% CAS-block overnight at 4°C. Tadpoles were washed 3 × 10 minutes at room temperature in PBS-Triton then blocked for 30 minutes in 10% CAS-block in PBS-Triton. Secondary antibody [goat anti-rabbit 488, Invitrogen A11008; goat anti-mouse 594, Abcam ab150116] was diluted 1:500 in 100% CAS-block and incubated for 2 hours at room temperature. Tadpoles were then washed 3 × 10 minutes in PBS-Triton followed by a 10 minute incubation in 1:2000 DAPI (Sigma D9542) in PBS-Triton before 3 × 20 minute washes in PBS-Triton. Isolated tails were mounted on slides in ProLong Diamond (ThermoFisher P36970).

### Light Microscopy

Images for Figure 1 were acquired on a Leica M205 stereo microscope with Leica DFC550 color camera using a Plan APO 1.0x objective. Images for Figures 2 and 3 were acquired on a Leica DM5500B upright microscope with motorized stage using 10x/0.30 HC PL Fluotar and 20x/0.50 HCX PL Fluotar objectives. Leica filter sets for GFP (Ex: 470/40 | Dc: 495 | Em: 525/50) and Texas Red (Ex: 560/50 | Dc: 585 | Em: 630/76) were used for immunofluorescent images. Images were collected using a 4 megapixel CCD sensor (Hamamatsu ORCA-Flash4.0 LT+) at 16-bit depth. Leica LAS X software was used for image acquisition and processing. All images were Extended Depth of Field processed from Z-stacks and fluorescent images of whole-mounted tadpole tails were created by stitching tiles from 3-4 acquisition regions.

### Quantification of Immunostaining results

For neurofilament bundle quantification, the number of neurofilament bundles present posterior of the vent were counted for stage 41 tadpoles injected with 1000 pg of WT or S160del hIMPDH2 mRNA, as well as uninjected controls.

For somitic boundary scoring, tadpoles injected with 1000 pg of WT or S160del hIMPDH2 mRNA, as well as uninjected controls, were assigned a somite boundary quality score by binning tails stained for skeletal muscle into 3 categories: well defined (normal appearance of somites), poorly defined (presence of abnormally-shaped somites or some lack of boundaries), and indistinguishable (no apparent somite boundaries).

### LC-MS sample preparation

For each aggregate tissue sample, 10 whole tadpoles were euthanized with a lethal dose of MS-222 at stage 41 and transferred into tubes. Media was removed, and samples were immediately flash frozen in liquid nitrogen. Five aggregate tissue samples were collected per group. Aqueous metabolites for targeted LC-MS profiling of 15 aggregate tissue samples were extracted using a protein precipitation method similar to the one previously described (34, 35) the Northwest Metabolomics Research Center.

Samples were first homogenized in 200 µL purified deionized water at 4 °C, and then 800 µL of cold methanol containing 124 µM 6C13-glucose and 25.9 µM 2C13-glutamate was added (reference internal standards were added to the samples in order to monitor sample prep). Afterwards samples were vortexed, stored for 30 minutes at −20 °C, sonicated in an ice bath for 10 minutes, centrifuged for 15 min at 14,000 rpm and 4 °C, and then 600 µL of supernatant was collected from each sample (left-over protein pallet was used for BCA assay). Lastly, recovered supernatants were dried on a SpeedVac and reconstituted in 0.5 mL of LC-matching solvent containing 17.8 µM 2C13-tyrosine and 39.2 3C13-lactate (reference internal standards were added to the reconstituting solvent in order to monitor LC-MS performance). Samples were transferred into LC vials and placed into a temperature controlled autosampler kept at 4 °C for LC-MS analysis.

### Targeted LC-MS Assay

Targeted LC-MS metabolite analysis was performed with a similar protocol as previously described by the Northwest Metabolomics Research Center on a duplex-LC-MS system composed of two Shimadzu UPLC pumps, CTC Analytics PAL HTC-xt temperature-controlled auto-sampler and AB Sciex 6500+ Triple Quadrupole MS equipped with ESI ionization source (35). UPLC pumps were connected to the auto-sampler in parallel and were able to perform two chromatography separations independently from each other. Each sample was injected twice on two identical analytical columns (Waters XBridge Premier BEH Amide column, Part # 186009930) performing separations in hydrophilic interaction liquid chromatography (HILIC) mode.

While one column was performing separation and MS data acquisition in ESI+ ionization mode, the other column was getting equilibrated for sample injection, chromatography separation and MS data acquisition in ESI-mode. Each chromatography separation was 16 minutes (total analysis time per sample was 32 minutes). MS data acquisition was performed in multiple-reaction-monitoring (MRM) mode. LC-MS system was controlled using AB Sciex Analyst 1.6.3 software. Measured MS peaks were integrated using AB Sciex MultiQuant 3.0.3 software. The LC-MS assay was targeting 373 metabolites (plus 4 spiked reference internal standards). Up to 175 metabolites (plus 4 spiked standards) were measured across the study set, and over 95% of measured metabolites were measured across all the samples. In addition to the study samples, two sets of quality control (QC) samples were used to monitor the assay performance as well as data reproducibility. One QC [QC(I)] was a pooled human serum sample used to monitor system performance. The other QC [QC(S)] was pooled study samples, and this QC was used to monitor data reproducibility. Each QC sample was injected per every 10 study samples. The data were highly reproducible with a median CV of 3.9 %. The data were normalized using the total protein count in each sample established with a BCA assay. Analysis of normalized data was done in MetaboAnalyst 6.0 and GraphPad Prism 10.

### MPA treatment

At two days post fertilization, tadpoles were transferred into 1/9x MR embryo rearing media supplemented with either 1 µM MPA or DMSO vehicle control and treated for 24 hours before fixation for immunostaining. Tadpoles were fixed in 1x MEM with 3.7% formaldehyde for 50 minutes at room temperature.

### Statistical analysis

For behavioral assays, percentages of tadpoles that swam and percentages of tadpoles that twitched were calculated, and a 95% confidence interval of each percentage was calculated using the Wilson/Brown method. Chi-square analysis indicated significant variation among groups, and the Marasculio procedure was used to compare WT and S160del, at P=0.05. Tail curve measurements were compared using one-way ANOVA and post hoc Tukey test for multiple comparisons. For metabolomics data, one-way ANOVA and post hoc Tukey test for multiple comparisons were used to compare metabolite abundances between groups. For neurofilament bundle quantification, one-way ANOVA and post hoc Tukey test for multiple comparisons were used to compare groups. For somitic boundary scoring, Fisher’s exact test was used to determine statistical significance. Analysis and graphing were done in GraphPad Prism 10.

### Recombinant protein expression and purification

WT and S160del hIMPDH2 protein was overexpressed in *E. coli* and purified as described previously (10). Briefly, hIMPDH2 with an N-terminal 6xHis-SMT3/SUMO tag was recombinantly expressed in BL21 (DE3) *E. coli* following induction with 1 mM IPTG for 4 hours at 30°C. Cells were lysed with an Emulsiflex, and His-tagged protein was isolated by affinity chromatography. The N-terminal tag was cleaved by incubation with ULP1, and protein was further purified by size exclusion chromatography on a Superose 6 Increase column, connected to an Äkta Pure FPLC system. Purified protein was concentrated to approximately 100 µM in Gel Filtration Buffer (20 mM HEPES, 100 mM KCl, 800 mM urea, and 1 mM DTT, pH 8) and flash-frozen in liquid nitrogen in single-use aliquots. Protein concentration was determined with a Bradford assay.

### Cryo-EM sample preparation

Purified S160del hIMPDH2 was incubated at a final concentration of 5 µM with 1 mM ATP, 1 mM IMP, 300 µM NAD^+^, and 20 mM GTP in Assay Buffer (20 mM HEPES, 100 mM KCl, and 1 mM DTT, pH 7). Samples were applied to glow-discharged C-flat holey carbon EM grids (Protochips), blotted, and plunge-frozen into liquid ethane using a Vitrobot (TFS) at 4°C and 100% humidity.

### Cryo-EM data collection and processing

High-throughput data collection was performed with SerialEM (36). Movies were collected in super-resolution mode on a Thermo Fisher Scientific Glacios TEM operating at 200 kV and equipped with a Gatan K3 Summit direct electron detector. Data collection parameters are summarized in Table 1.

**Table 1.** Cryo-EM data collection and model refinement statistics.

|  | **hIMPDH2-S160del**<br>**Interfacial octamer** | **hIMPDH2-S160del**<br>**Tetramer** |
| --- | --- | --- |
| **Ligands** | GTP, ATP, IMP, NAD<sup>+</sup> | GTP, ATP, IMP, NAD<sup>+</sup> |
| **PDB ID** | 9MUC | 9MUB |
| **EMDB ID** | EMD-48628 | EMD-48627 |
| **Data collection and refinement** |  |  |
| **Magnification** | 45,000 | 45,000 |
| **Voltage (kV)** | 200 | 200 |
| **Electron exposure (e<sup>-</sup>/Å<sup>2</sup>)** | 50 | 50 |
| **Nominal defocus range (μm)** | -1.8 — -1.2 | -1.8 — -1.2 |
| **Pixel size (data collection) (Å)** | 0.4425 | 0.4425 |
| **Pixel size (reconstruction) (Å)** | 0.885 | 0.885 |
| **Micrographs (no.)** | 2,630 | 2,630 |
| **Initial particles (no.)** | 1,393,912 | 1,393,912 |
| **Final particles (no.)** | 615,147 | 259,441 |
| **Symmetry imposed** | D4 | C4 |
| **Resolution, CryoSPARC**<br>**postprocess (0.143 FSC) (Å)** | 2.1 | 2.5 |
| **Resolution, Phenix density**<br>**modification (0.5 FSC<sub>ref</sub>) (Å)** | 2.1 | 2.3 |
| **Model refinement and validation** |  |  |
| **Initial model (PDB ID)** | 6UA5 | 6UA5 |
| **R.m.s. deviations** |  |  |
| **Bond lengths (Å)** | 0.0162 | 0.0162 |
| **Bond angles (°)** | 1.90 | 1.99 |
| **MolProbity score** | 1.17 | 1.02 |
| **Clashscore** | 1.33 | 1.35 |
| **C-beta deviations** | 0.00% | 0.62% |
| **Rotamer outliers (%)** | 2.57% | 1.49% |
| **Ramachandran plot** |  |  |
| **Favored (%)** | 98.20% | 97.88% |
| **Allowed (%)** | 1.80% | 2.12% |
| **Disallowed (%)** | 0.00% | 0.00% |

During data collection, CryoSPARC Live was used for initial steps of data processing (37). Movies were aligned with Patch Motion Correction with an output Fourier cropping factor of 0.5. Contrast transfer function estimation was performed with Patch CTF Estimation. Blob picking was done with a particle diameter set to 200 Å. Particles were extracted with a box size of 300 pixels and Fourier cropped to 150 pixels. 2D classification and 2D class selection were done in CryoSPARC Live. A final selection of particles and micrographs were exported from CryoSPARC Live, and particles were re-extracted from micrographs without Fourier cropping. Downstream processing to sort particles into interfacial octamer, tetramer, and canonical octamer classes was done in CryoSPARC v3 or v4.

Four Ab-initio Reconstructions were generated from unbinned particles, with a maximum resolution of 20 Å. One class resembled an interfacial octamer. Homogenous refinement of this class with C1 symmetry, using the Ab-initio map as the initial volume, resulted in a volume which appeared D4-symmetric. Iterative rounds of 3D classification to purify the interfacial octamer and Non-Uniform Refinement with D4 symmetry resulted in a high resolution map of the interfacial octamer, to be used as an initial volume in downstream processing.

The other 3 classes from the Ab-initio Reconstruction job were fed into Heterogenous Refinement with 2 classes, using a low-pass filtered extended octamer Molmap generated in UCSF Chimera as the initial model for each class. The class most resembling a tetramer was selected, and a second round of Heterogeneous Refinement was performed to further purify out the tetramer. Iterative rounds of 3D classification to remove heterogeneity, followed by Non-Uniform Refinement with C4 symmetry, resulted in a high resolution map of the tetramer, to be used as an initial volume in downstream processing.

These volumes of the interfacial octamer and tetramer, as well as an imported volume of a canonical IMPDH octamer (EMD-20701), were used as initial volumes in a Heterogeneous Refinement job on the original 920,210 particles, to sort the entire dataset into interfacial octamer, tetramer, and canonical octamer classes. For the interfacial octamer and tetramer classes, per-particle defocus refinement, and CTF refinement, followed by Non-Uniform refinement, with D4 or C4 symmetry imposed, improved resolution further. PHENIX v1.20.1-4487 Density Modification improved the interpretability of the volumes for model building.

### Model building and refinement

For the interfacial octamer, the structure of the WT hIMPDH2 free interfacial octamer was used as an initial model (PDB: 6UA5). The initial model was rigid-body fit into the final volume in UCSF ChimeraX v1.6.1 (38), and automated fitting was done with real space refinement in PHENIX v1.20.1-4487, with rigid-body refinement, noncrystallographic symmetry constraints, gradient-driven minimization, and simulated annealing (39, 40). The output was manually adjusted residue-by-residue in Coot v0.9.8.8 (41) and with semi-automated fitting in Isolde v1.6.0 (42). These steps were repeated iteratively to improve fit and Molprobity statistics. The final model of the interfacial octamer was used as an initial model for the tetramer, and the same model refinement process was used. Refinement statistics for the interfacial octamer and tetramer structures are reported in Table 1. Figures were prepared using UCSF ChimeraX.

### Data availability

Coordinates for cryo-EM structures and maps are deposited in the Protein Data Bank and Electron Microscopy Data Bank, respectively, with the following accession IDs: 9MUC and EMD-48628 for the S160del interfacial octamer; 9MUB and EMD-48627 for the S160del tetramer.

The metabolomics dataset was deposited at metabolomicsworkbench.org under the Study ID ST003858. Any additional information required to re-analyze the data will be made available by the corresponding author upon request.

### Negative stain electron microscopy

Purified protein was diluted to a final concentration of 1 µM in Assay Buffer (20 mM HEPES, 100 mM KCl, 1 mM DTT, pH 7.0) and incubated with Ap5G, GTP, IMP, and NAD^+^ for 10 minutes at room temperature. Sample was applied to glow-discharged continuous carbon EM grids and negatively stained with 2% (w/v) uranyl formate. Grids were imaged on a 120 kV FEI Tecnai Spirit microscope with a 4k x 4k Gatan Ultrascan CCD camera.

### *In vitro* enzyme assays

Enzyme assays were conducted as previously described (4, 10). Briefly, purified protein was diluted to a final concentration of 1 µM in Assay Buffer (20 mM HEPES, 100 mM KCl, 1 mM DTT, pH 7.0) and pre-incubated with Ap5G, GTP, and IMP for 15 minutes at 25°C. The reaction was initiated with the addition of NAD^+^. The final concentration of ligands used in the assay was 1 mM Ap5G, 5 mM GTP, 1 mM IMP, and 300 µM NAD^+^. Absorbance at 340 nm was measured over time in increments of 1 minute, using a Varioskan Lux microplate reader (Thermo Fisher Scientific). NADH production was calculated from absorbance using a standard curve, and specific activity was calculated by the slope for a 4 min window. All data points are an average of three measurements from the same protein preparation. Error bars represent standard deviation.

## ACKNOWLEDGEMENTS

We thank the Arnold and Mabel Beckman Cryo-EM Center at the University of Washington for electron microscope use. We also thank the Northwest Metabolomics Research Center for performing LC-MS. This work was supported by a University of Washington Levinson fellowship to M.E.M. and by the US National Institutes of Health (R35GM149542 to J.M.K., 5R01GM148490 and 5R01NS099124 to A.E.W., and T32GM008268 to A.G.O.).

---

**Supplemental Figure 1.**
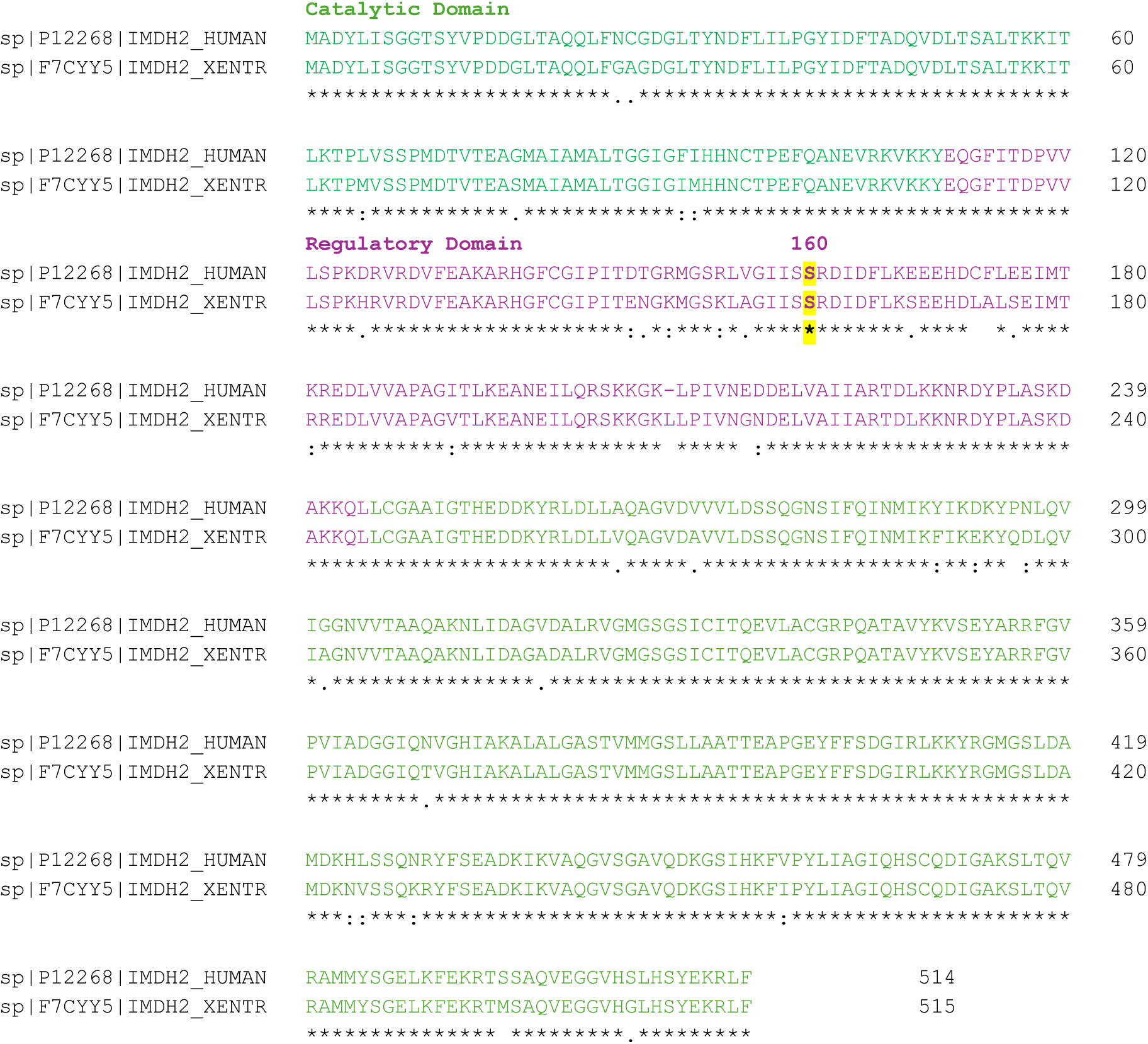
**IMPDH2 sequence alignment** Protein sequence alignment of human and *Xenopus tropicalis* IMPDH2 generated with Clustal O (1.2.4) MSA. Green color represents the catalytic domain and purple color represents the regulatory Bateman domain. The conserved serine residue 160 is highlighted.

**Supplemental Figure 2.**
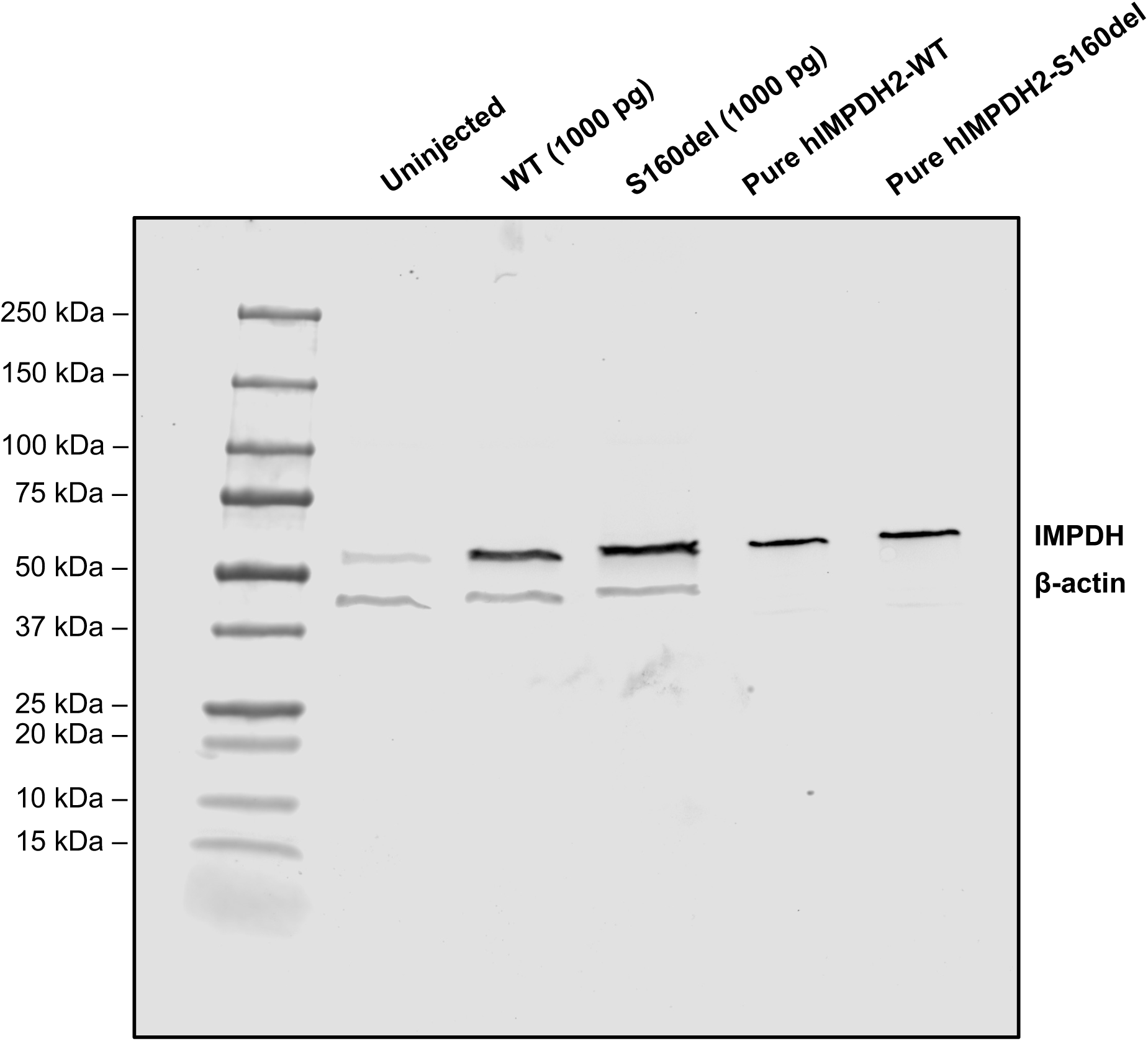
**Uncropped western blot of injected embryos at stage 21** Uncropped Western blot of injected Xenopus embryos at NF stage 21, approximately 1 dpf, shown in Figure 1D.

**Supplemental Figure 3.**
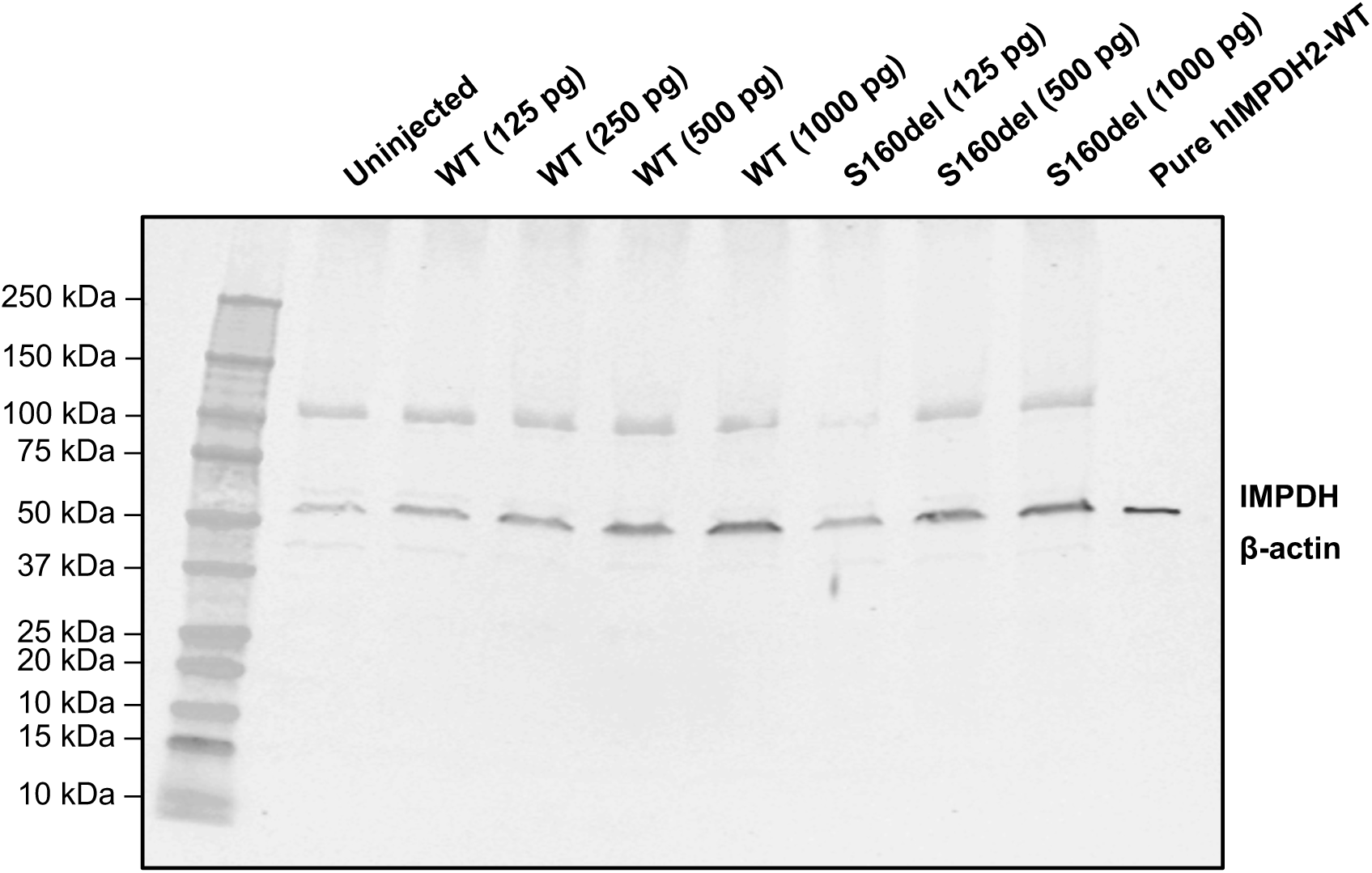
**Dose-dependent overexpression of hIMPDH2 mRNA** Representative western blot analysis of the mRNA dose response on hIMPDH2 protein expression in Xenopus embryos at NF stage 21, approximately 1 dpf. Trend was seen across two independent experiments.

**Supplemental Figure 4.**
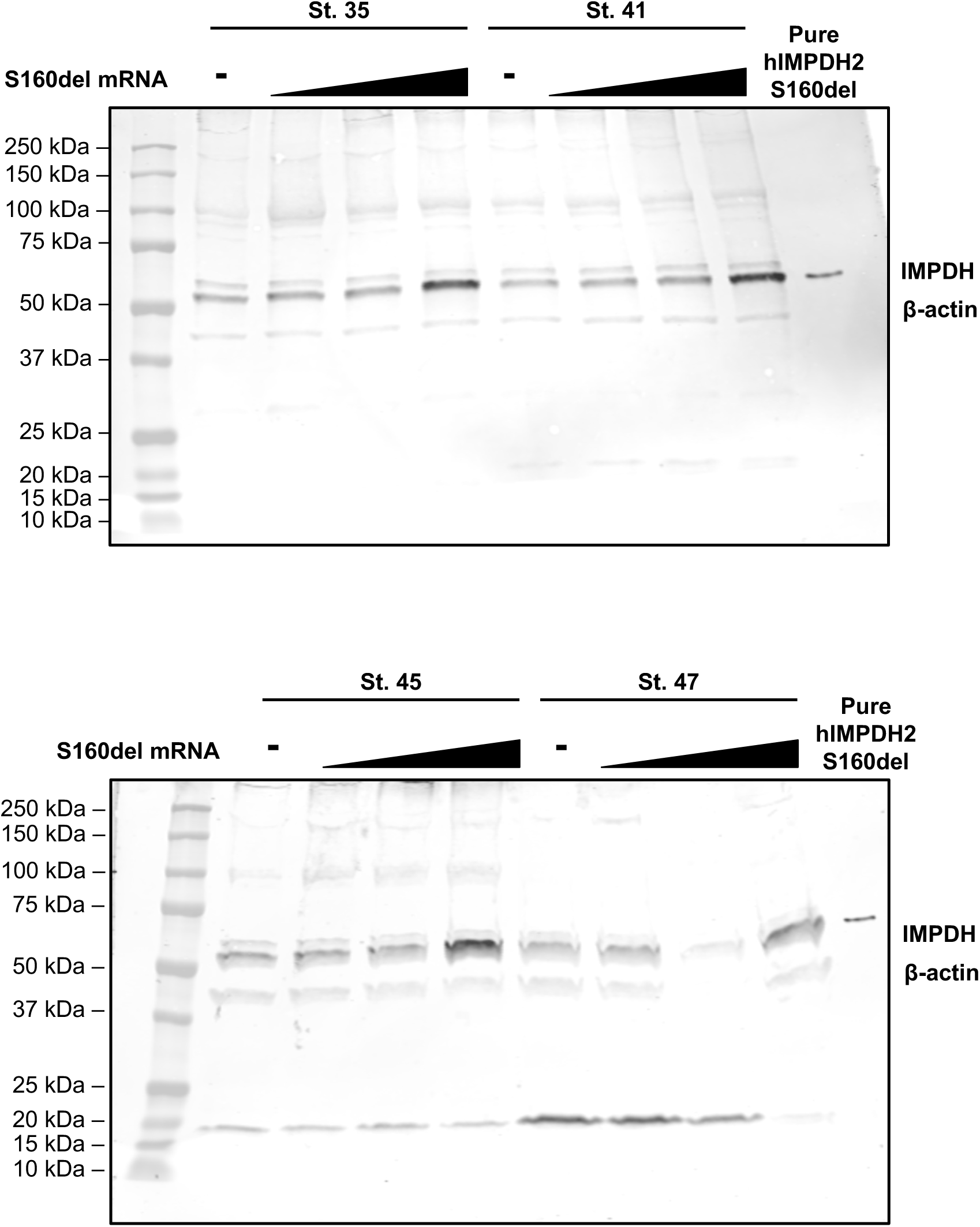
**Time-course western blot of tadpoles injected with S160del mRNA** Western blot analysis of S160del hIMPDH2 expression in injected tadpoles from NF stages 35-47, or approximately 2-5 dpf. The doses of mRNA injected were 0 pg, 250 pg, 500 pg, and 1000 pg.

**Supplemental Figure 5.**
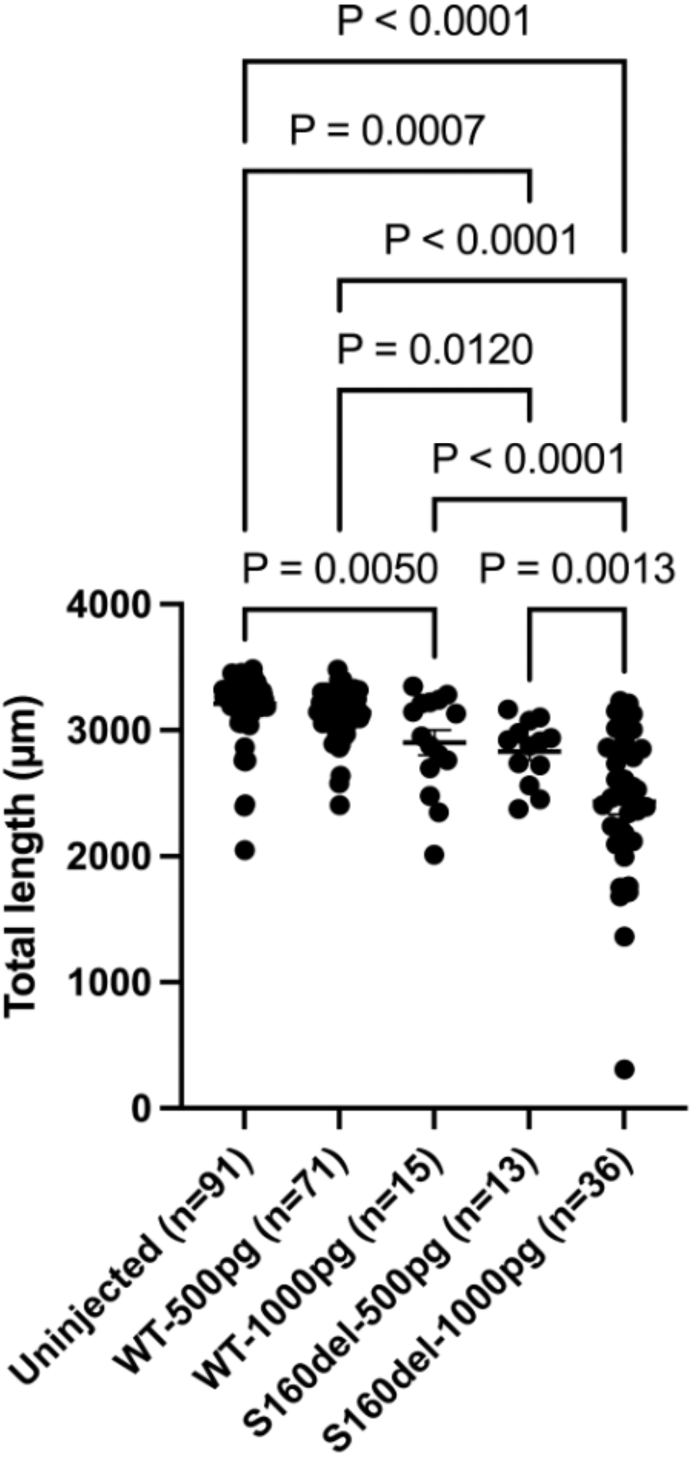
**Length of tadpoles at stage 41** Mean length of tadpoles at stage 41, approximately 3 dpf, measured from nose to tail tip, along the spinal cord. Error bars represent the SEM.

**Supplemental Figure 6.**
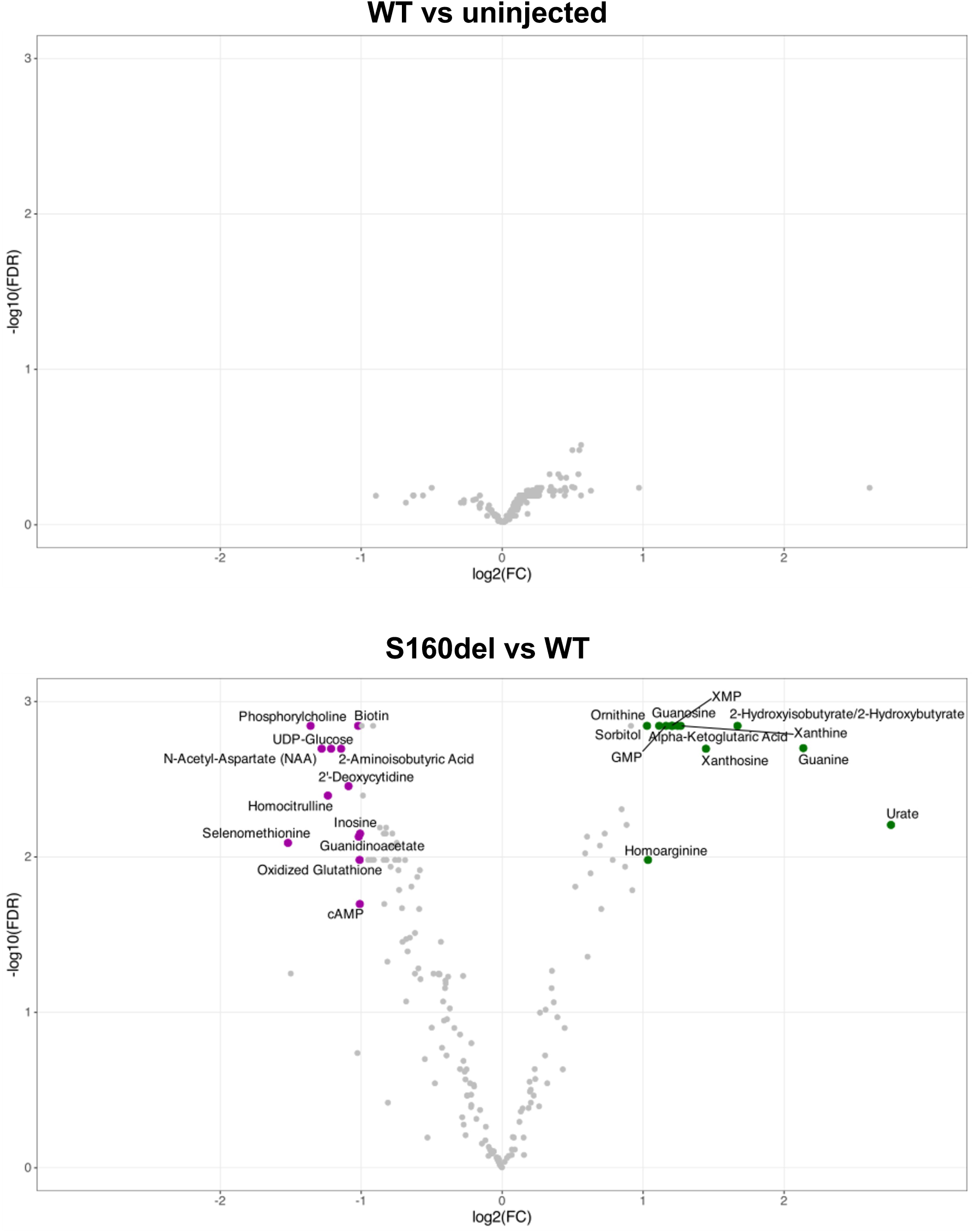
**Volcano plots of metabolomics data** Volcano plots of metabolite levels comparing WT hIMPDH2 expressing to uninjected tadpoles (A) and comparing S160del hIMPDH2 expressing to WT hIMPDH2 expressing tadpoles (B). Labels represent metabolites with a Bonferroni false discovery rate of less than 0.05 and a fold-change of at least 2.

**Supplemental Figure 7.**
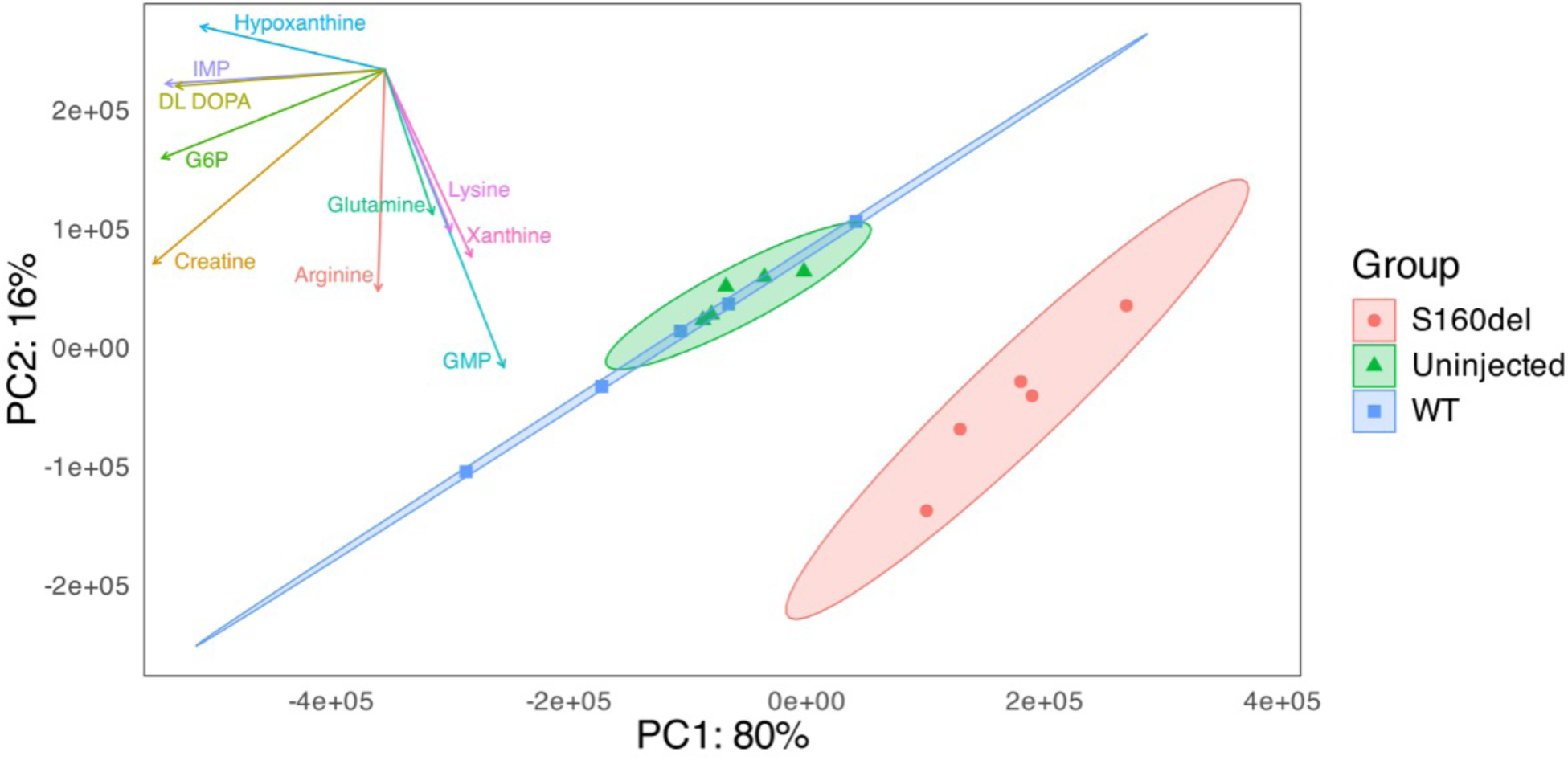
**Principal component analysis of metabolomics dataset** Principal component analysis of metabolomics data. Loading vectors for the 10 most weighted metabolites in PC1 are shown in the top-left corner.

**Supplemental Figure 8.**
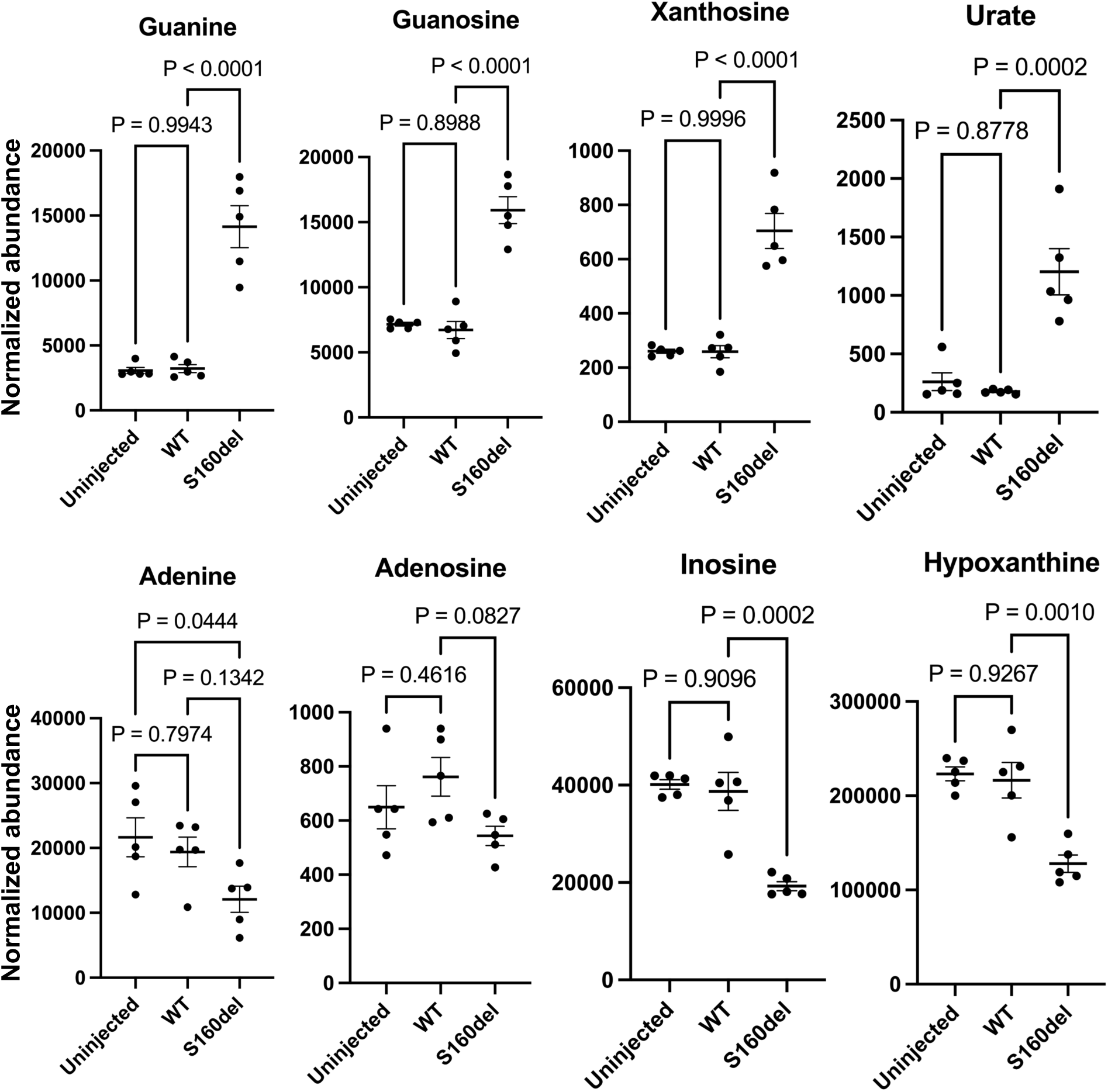
**Other metabolites of interest** Additional plots of other metabolite levels relevant to purine synthesis and degradation. Each data point represents normalized metabolite abundance of an aggregate tissue sample composed of 10 whole tadpoles. One-way ANOVA and Tukey’s multiple comparisons test performed on metabolite levels normalized to total protein from BCA. Plotting mean +/- SEM.

**Supplemental Figure 9.**
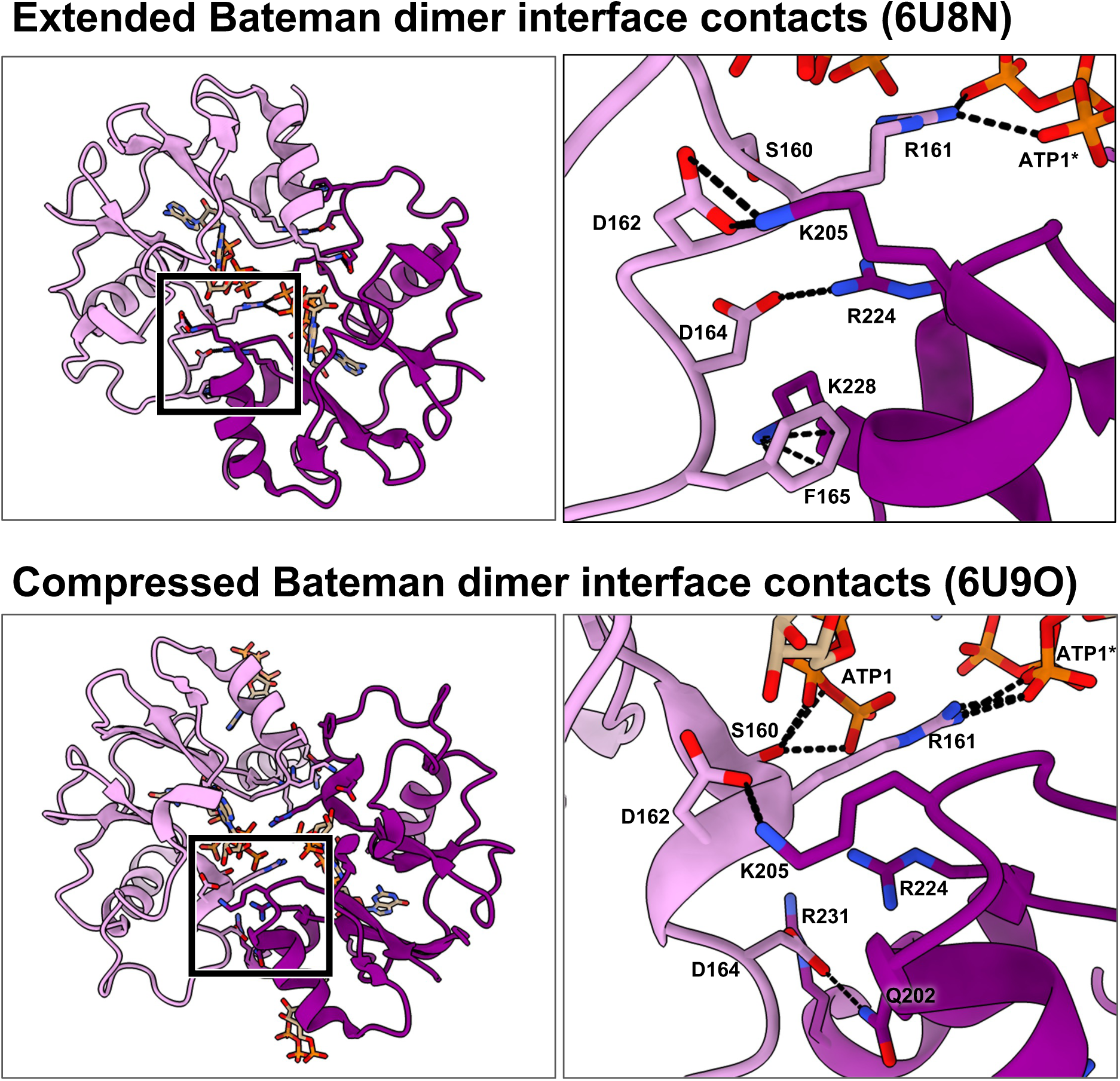
**Bateman dimer interface contacts in WT hIMPDH2** Contacts at the Bateman dimer interface in previously published structures of extended and compressed WT human IMPDH2 filaments.

**Supplemental Figure 10.**
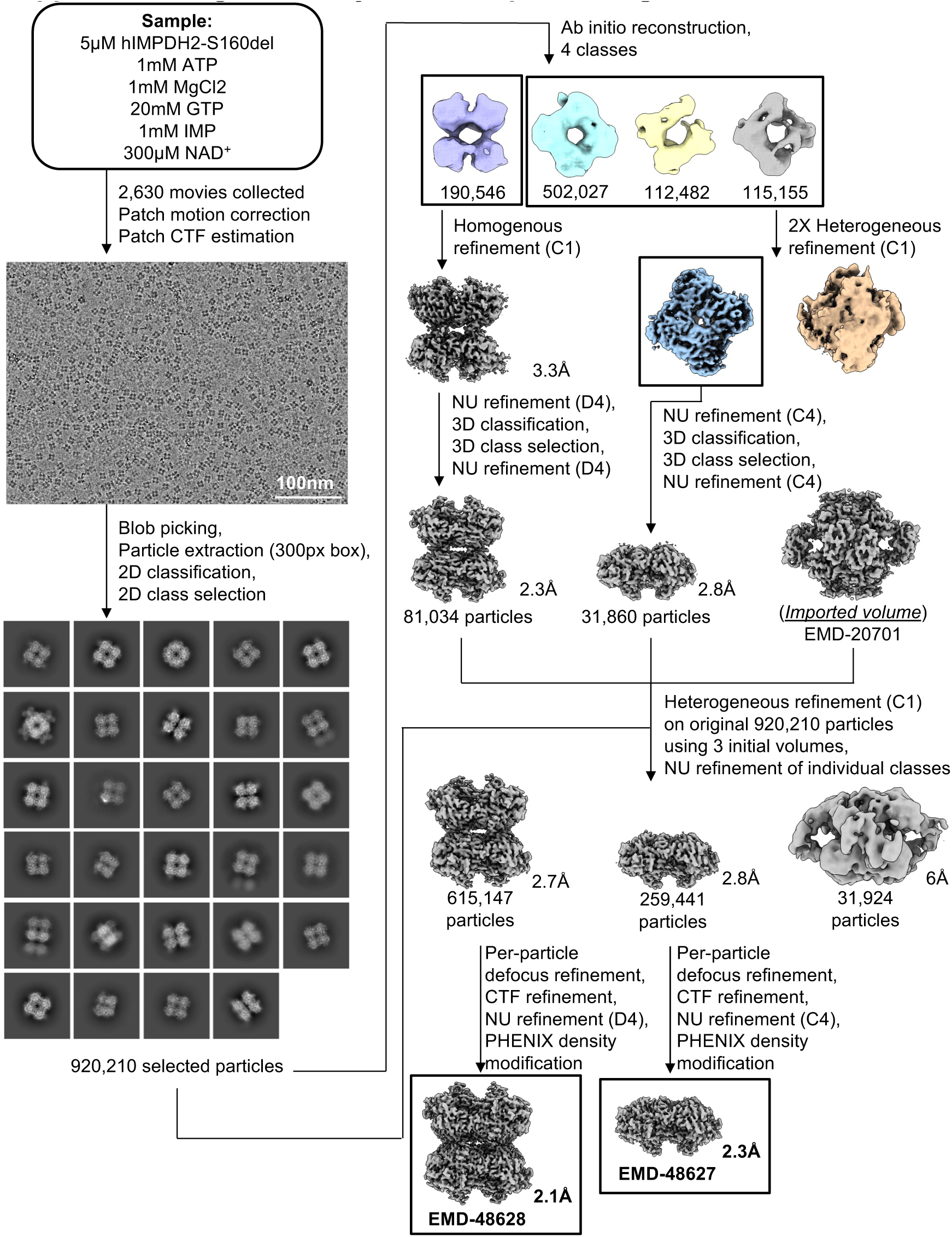
**Cryo-EM data processing** Cryo-EM data processing workflow to classify between interfacial octamers, tetramers, and canonical octamers.

**Supplemental Figure 11.**
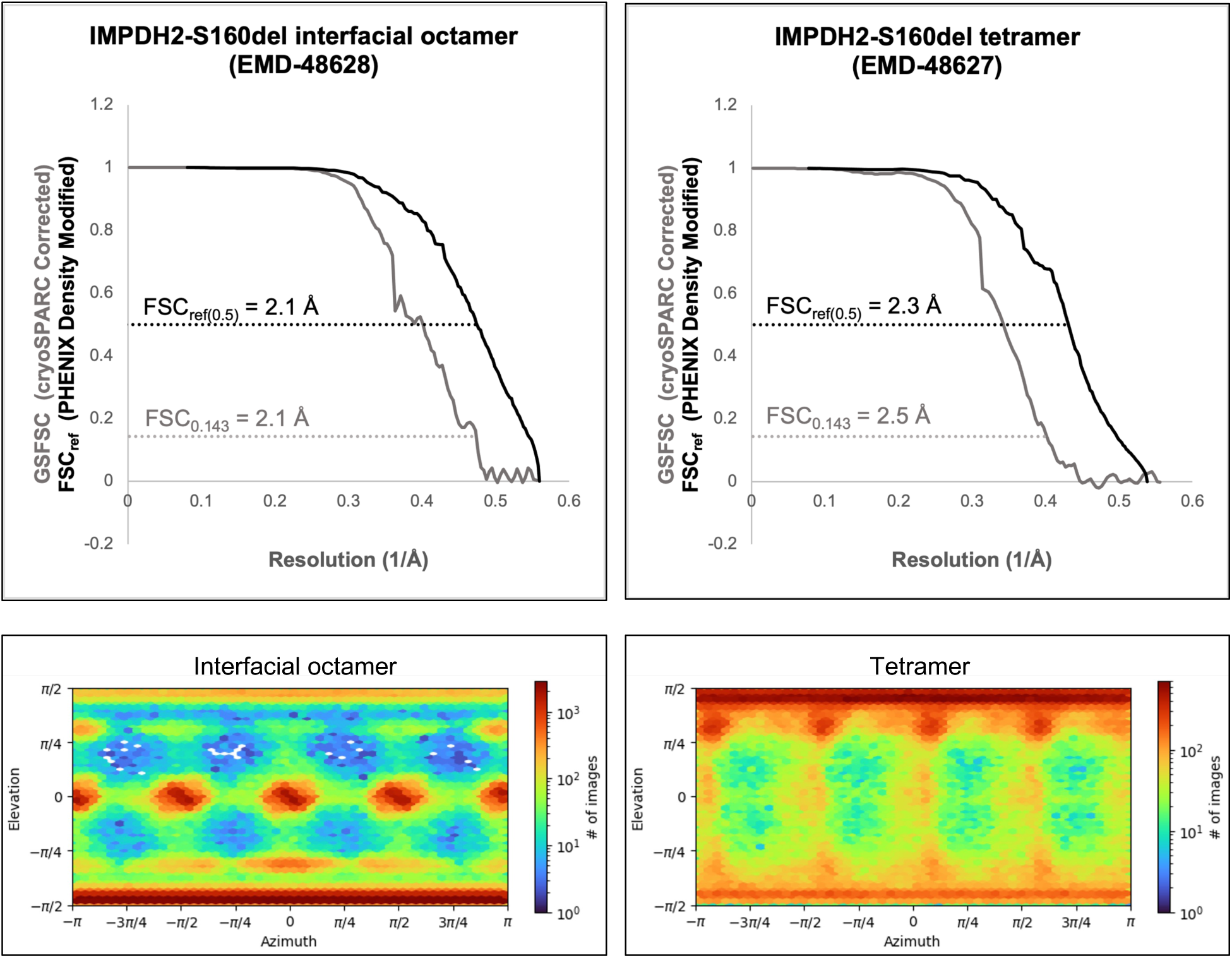
**FSC curves and viewing direction distributions of cryo-EM structures** FSC curves and viewing distributions of the interfacial octamer and tetramer S160del reconstructions.

**Supplemental Figure 12.**
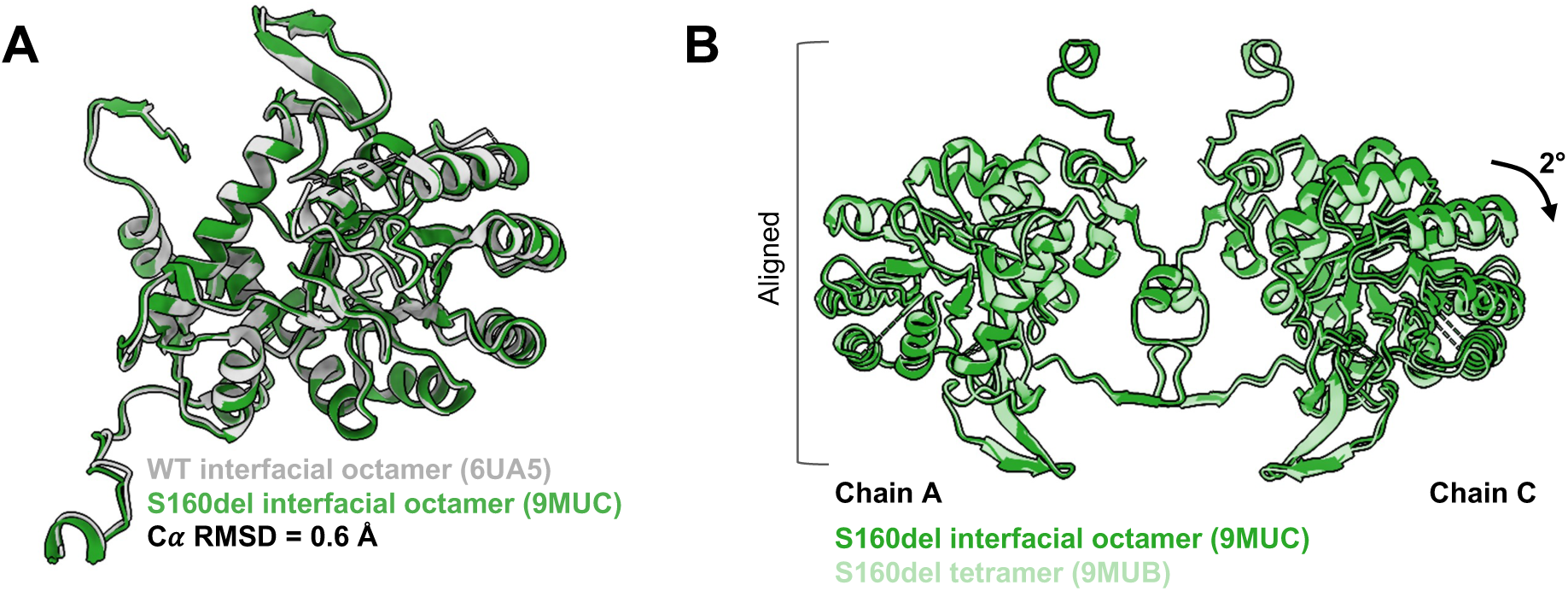
**Cryo-EM model comparisons** A. The S160del interfacial octamer structure (green) aligned to the previously published WT hIMPDH2 interfacial octamer structure (gray). Bateman domains are highly mobile and unresolved in both structures. The alpha-carbon RMSD of 0.6 Å suggests no major differences in the catalytic domain. B. The S160del interfacial octamer aligned to the S160del tetramer at Chain A. The filament assembly interface of the interfacial octamer is flat compared to the tetramer, which is bowed by 2 degrees.

**Supplemental Figure 13.**
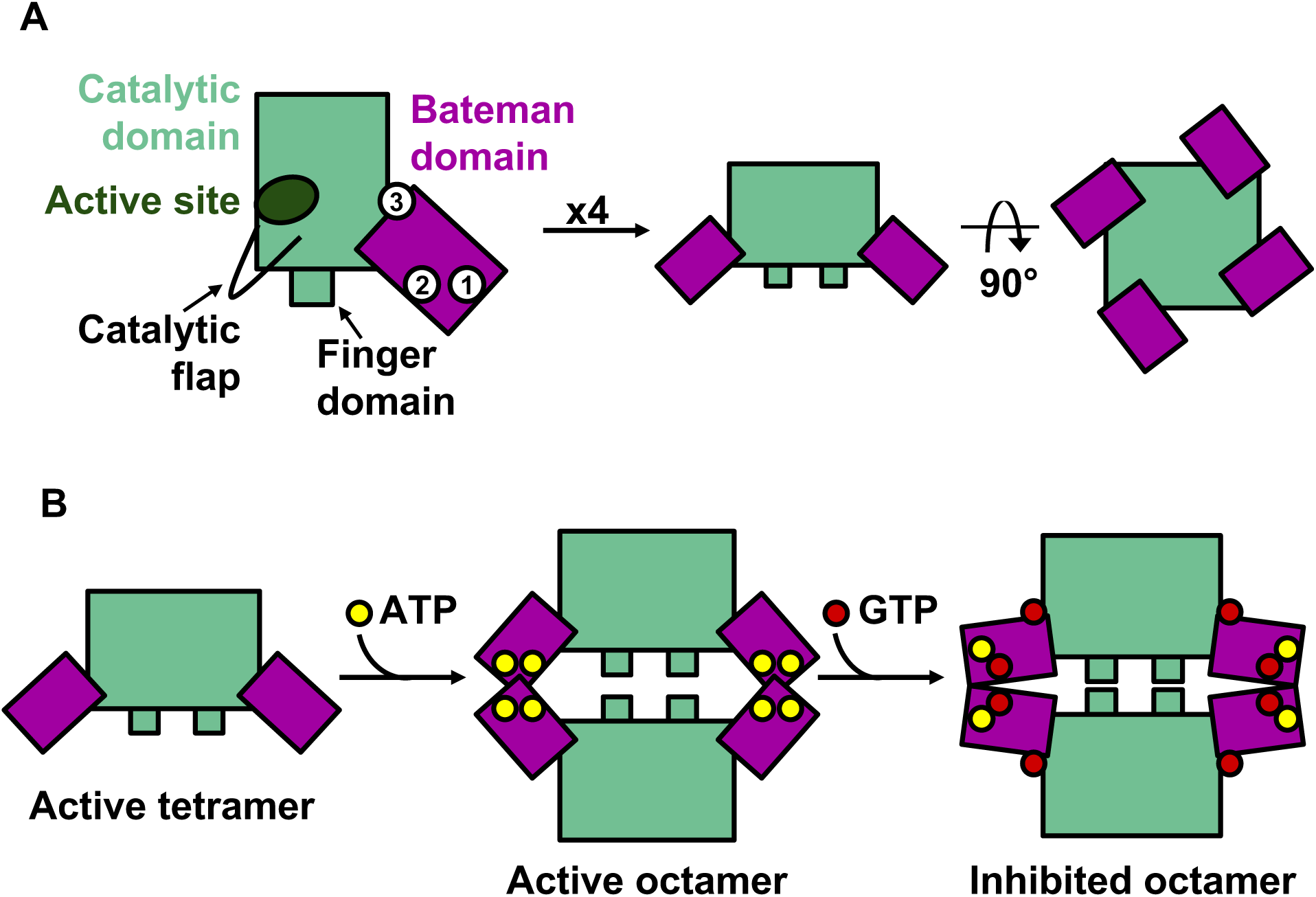
**IMPDH structure and assembly** A. IMPDH is composed of the catalytic domain (green) and the regulatory Bateman domain (purple), which contains three allosteric sites where nucleotides can bind. The catalytic domain contains the active site, where IMP and NAD^+^ bind (dark green), the catalytic flap, and the finger domain. Tetramers form through interactions between catalytic domains. B. Upon ATP binding to allosteric sites 1 and 2, IMPDH tetramers dimerize through interactions between Bateman domains. These octamers are extended and catalytically active. GTP binding to allosteric sites 2 and 3 induces compression of the octamer and interaction between finger domains of opposing tetramers, resulting in inhibition.

**Supplemental Movie 1.**

Behavior of a stage 41 uninjected tadpole.

**Supplemental Movie 2.**

Behavior of a stage 41 tadpole injected with 1000 pg of hIMPDH2-WT mRNA.

**Supplemental Movie 3.**

Behavior of a stage 41 tadpole injected with 1000 pg of hIMPDH2-S160del mRNA.

**Supplemental Movie 4.**

Group of four stage 41 tadpoles injected with 1000 pg of hIMPDH2-S160del mRNA exhibiting twitching behavior.

